# A multiplex platform to identify mechanisms and modulators of proteotoxicity in neurodegeneration

**DOI:** 10.1101/2022.09.19.508444

**Authors:** Samuel J. Resnick, Seema Qamar, Jenny Sheng, Lei Haley Huang, Jonathon Nixon-Abell, Schuyler Melore, Chyi Wei Chung, Xuecong Li, Jingshu Wang, Nancy Zhang, Neil A. Shneider, Clemens F. Kaminski, Francesco Simone Ruggeri, Gabriele S. Kaminski Schierle, Peter St George-Hyslop, Alejandro Chavez

## Abstract

Neurodegenerative disorders are a family of diseases that remain poorly treated despite their growing global health burden. A shared feature of many neurodegenerative disorders is the accumulation of toxic misfolded proteins. To gain insight into the mechanisms and modulators of protein misfolding, we developed a multiplex reverse genetics platform. Using this novel platform 29 cell-based models expressing proteins that undergo misfolding in neurodegeneration were probed against more than a thousand genetic modifiers. The resulting data provide insight into the nature of modifiers that act on multiple misfolded proteins as compared to those that show activity on only one. To illustrate the utility of this platform, we extensively characterized a potent hit from our screens, the human chaperone DNAJB6. We show that DNAJB6 is a general modifier of the toxicity and solubility of multiple amyotrophic lateral sclerosis and frontotemporal dementia (ALS/FTD)-linked RNA-binding proteins (RBPs), including FUS, TDP-43, and hnRNPA1. Biophysical examination of DNAJB6 demonstrated that it co-phase separates with, and alters the behavior of FUS containing condensates by locking them into a loose gel-like state which prevents their fibrilization. Domain mapping and a deep mutational scan of DNAJB6 support the critical importance for DNAJB6 phase separation in its effects on multiple RNA-binding proteins. Crucially, these studies also suggest that this property can be further tuned to generate novel variants with enhanced activity that might illuminate potential avenues for clinical translation.

## Introduction

Human neurodegenerative diseases are a major source of morbidity and mortality worldwide and represent a significant unmet medical need^1–3^. A hallmark of many neurodegenerative diseases (NDDs) is the intracellular accumulation of misfolded protein aggregates^4,5^. To model the proteotoxicity imposed by NDD-associated proteins with an intrinsic propensity to aggregate such as Fused in Sarcoma (FUS), TAR DNA-binding protein (TDP-43), and alpha-synuclein, researchers have repeatedly turned to the yeast, *Saccharomyces cerevisiae*^6–8^. In yeast, expression of these aggregation-prone proteins results in slow growth. By screening for genes that restore growth upon overexpression, researchers have been able to identify pathways involved in modulating the underlying proteotoxicity, and validate their findings in lower throughput mammalian cell systems. As a whole, these screens have informed our understanding of disease pathobiology, and served as the basis for numerous therapeutic intervention strategies, some of which are being advanced by commercial entities^7–18^.

While previous yeast-based studies have proven insightful, only a subset of NDD models have been screened. This likely reflects a systems-level limitation imposed by the significant effort involved in conducting large genetic screens. In addition, due to differences in the testing environment, genetic background, and method of screening, it is difficult to definitively compare results across studies in order to gain insight into broad versus narrow-acting regulators of proteotoxicity^7,8,18^. When comparisons between screens have been performed, few or no shared toxicity modifiers have been identified, even among related disease proteins such as TDP-43 and FUS^7,8,10^. This lack of correlation conflicts with overlaps in clinical presentation between TDP-43 and FUS patients and the shared biochemical and biophysical properties of both proteins^19^. Furthermore, as previous approaches are only able to study one model at a time, they have mainly focused on screening wild-type versions of NDD associated proteins, preventing our ability to assess if patient mutations alter the underlying molecular processes, which if found, would have significant implications in the treatment of this family of disorders^8,15^. Finally, most screens performed in yeast have searched for yeast genes that rescue the toxicity of NDD models, leading to identified hits with unclear or no known orthologous human counterpart to advance as a potential therapeutic candidate^8,10^.

To overcome these limitations, we created a multiplex screening platform capable of simultaneously identifying genetic suppressors to 29 cell-based models expressing proteins that undergo misfolding in neurodegeneration. To interpret the wealth of data obtained, we built a custom analysis pipeline that prioritizes interactions for subsequent validation. Using this platform, we were able to identify previously elusive broadly-active rescuers, along with highly selective rescuers that only impact a single model. These studies revealed a plethora of genetic modifiers for future investigation, along with highlighting the diverse array of pathways and mechanisms that can potentially be exploited for therapeutic benefit.

Upon further examination of our results, we identified the human HSP40 co-chaperone, DNAJB6, as a potent rescuer of the toxicity caused by the expression of multiple RNA-binding proteins associated with ALS/FTD. Through subsequent studies in mammalian cells, we show that DNAJB6 has the ability to modulate the solubility of FUS, TDP-43, and heterogeneous nuclear ribonucleoprotein A1 (hnRNPA1). We also use purified proteins to demonstrate that DNAJB6 is able to phase separate and alter the liquid-liquid phase separation properties of FUS. We show that DNAJB6 is able to maintain FUS in a loose gel-like state that prevents its fibrilization over extending time periods. This mechanism is unique among modifiers of biologic condensates and suggests an additional mechanism by which chaperones prevent the aggregation of clients. We corroborate our *in vitro* findings by analyzing a series of truncation mutants and performing a deep mutational scan within DNAJB6, along with suggesting mechanisms by which its activity can be further engineered.

## Results

### Development of a multiplexed screening strategy to identify rescuers of proteotoxicity

To enable our multiplex screening approach, we make use of isogenic yeast strains that each contain a unique DNA-barcode inserted into a neutral genomic locus^20^. Into each of these barcoded strains, we deliver a construct encoding a NDD-associated protein with a propensity to misfold (e.g., TDP-43, FUS, alpha-synuclein). Growth of these strains in media that induces the overexpression of the toxic disease-associated protein causes a reduction in cell growth, and provides a facile method of modeling the cellular dysfunction elicited by these proteins. As each of our models (i.e. yeast expressing a unique protein of interest) are linked to a particular DNA-barcode, we can combine them into a single mixed pool and track the growth of each member by measuring its barcode abundance using next-generation sequencing. To identify novel regulators of neurodegeneration, we probe the pool of disease models against a library of genetic modifiers (Sup. Fig. 1). In cases where a genetic modifier suppresses the toxicity of a particular NDD-associated protein, we observe a marked increase in the abundance of the model’s barcode as compared to the control condition where the pool is exposed to an inert genetic modifier like mCherry. Taking advantage of the scalability afforded by the use of DNA-barcoding, each examined protein is placed into several different DNA-barcoded strains (i.e. redundant barcoding). This decreases assay noise by allowing us to use the collective behaviors of all uniquely barcoded strains containing the same protein to derive our conclusions and enables us to confidently identify significant interactions between our disease models and genetic modifiers^21^.

To develop our approach, we analyzed multiple screening parameters such as strategies for pooling DNA-barcoded strains based on their toxicity, appropriate number of experimental replicates, and the amount of redundant barcoding required to detect interactions with high sensitivity (Sup. Note 1-3, Sup. Fig. 1-7). We then assembled a collection of NDD-associated proteins based on previous publications (Fig. 1a and Sup. Table 1)^13,22–24^. For a subset of these proteins, we also engineered a panel of point mutants based on familial variants that increase the likelihood of disease to determine whether they might influence the observed rescue (Fig. 1a). To assist in interpreting screening results, the final pool of cells was supplemented with a series DNA-barcoded controls including cells that express a non-toxic fluorescent protein (mCherry), other aggregation-prone proteins not associated with neurodegeneration (e.g. SUP35, RNQ1), or proteins linked to neurodegeneration but which themselves are not prone to misfolding (e.g. ANG, OPTN). In total, our final screening library contains 29 NDD-associated proteins plus multiple controls, each placed within 5-7 uniquely DNA-barcoded strains all mixed together into a single pool for testing.

**Figure 1.**
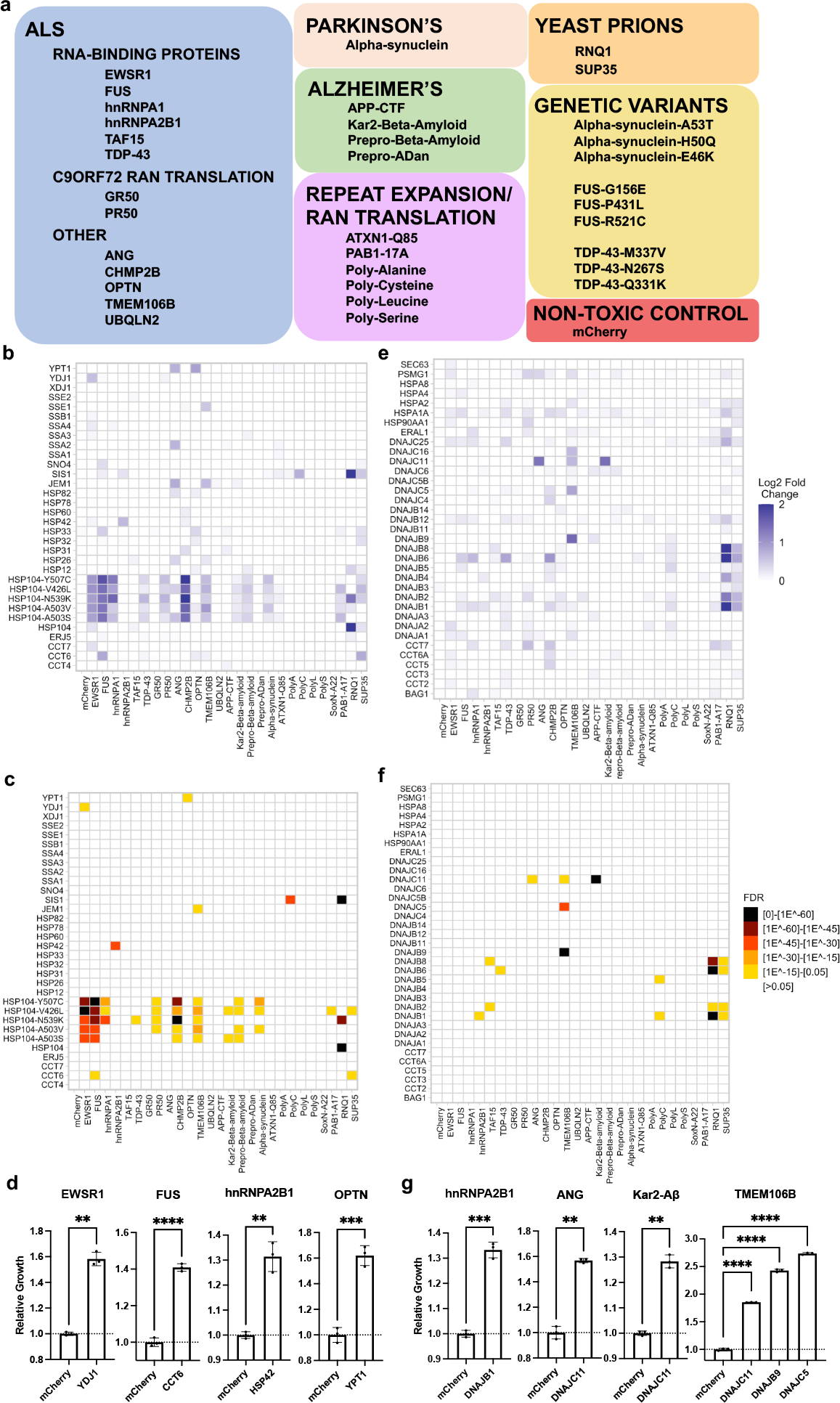
Screening of molecular chaperones from yeast and humans for their ability to rescue the proteotoxicity of various neurodegenerative disease and protein misfolding models. **a**. Models included in screen and disease association. **b**. Log2 fold change plotted for interactions between selected yeast chaperones and models. Log2 fold changes are shown as an average of all barcoded strains associated with that model **c**. Statistically significant interactions between selected yeast chaperones and models. **d**. Validation of interactions between yeast chaperones and models. Data are shown as mean ± s.d. for three biological replicates. **e**. Log2 fold change plotted for interactions between selected human chaperones and models **f**. Statistically significant interactions between selected human chaperones and models. **g**. Validation of interactions between human chaperones and models. Data are shown as mean ± s.d. for three biological replicates. Comparisons between two conditions were conducted with Welch’s t tests while multiple comparisons were conducted with ordinary one-way ANOVA; **P≤0.01, ***P≤0.001,****P≤0.0001.

### Functional characterization of chaperone interactions with neurodegenerative disease models

Using our final pool and optimized screening strategy, we probed a targeted library of 132 molecular chaperones, 62 of which were from yeast and 70 from humans. Molecular chaperones have been previously implicated in the refolding, turnover, and mitigation of the toxicity of aggregation-prone proteins associated with neurodegeneration^5,25,26^. However, a comprehensive map of the functional interactions between chaperones and their disease-associated clients remains elusive, limiting the field’s ability to identify broadly-active members of this class of proteins. We hypothesized that the approach described here would be uniquely suited towards assessing this interaction space. In addition, it would also serve as a data-rich set to extensively validate our methodology. Overall, this screen represents 5,850 genetic interactions between the various proteotoxic models and controls and the corresponding library of molecular chaperones.

Within the yeast chaperone set, we observed 112 strong interactions that resulted in a greater than 0.5 log2 fold increase in barcode abundance (Fig. 1b, Sup. Fig. 8a, and Sup. Table 2). Notably, potentiated HSP104 chaperones were broadly active with the ability to rescue the proteotoxicity of a number of models including two beta-amyloid models, hnRNPA1, and the dipeptide repeat PR50 associated with C9orf72 RAN-translation, in addition to their previously reported rescue of TDP-43, FUS, and alpha-synuclein models^27^. Outside of strong interactions, such as those observed with the HSP104 variants, 74 interactions with mild-to-moderate positive log2 fold changes between 0.25 and 0.5 were also observed. In order to prioritize the 186 interactions for further follow up, we developed an analysis pipeline to call interactions with statistically significant enrichment (Fig. 1c, and Sup. Fig. 8b). This pipeline uses the control wells tested against inert rescuers (e.g. mCherry) to develop an expectation for the abundance of each barcode in the pool, it then determines which barcodes significantly increase in abundance in test wells, and combines information from barcodes associated with the same model to identify the most potent and significant hits (see Materials and Methods for details). Upon applying our data analysis pipeline, 100 interactions were called as significant (Sup. Table 2). Among these significant interactions, we identified specific interactions between the ALS/FTD-associated RNA-binding proteins, EWSR1, FUS, and hnRNAP2B1 and the type I HSP40 chaperone YDJ1, the chaperonin containing TCP-1 (CCT) subunit CCT6, and the small heat shock protein HSP42, respectively. We also observed an interaction between the Golgi-maintenance protein and autophagosomal receptor OPTN and a Rab family GTPase involved with ER-to-Golgi transport that localizes to pre-autophagosomal structures, YPT1 (Fig. 1d).

The mammalian chaperone set contained 131 interactions that resulted in mild-to-moderate or strong log2 fold changes in barcode abundance (Fig. 1e and Sup. Fig. 9). However, 86 of the 131 interactions led to mild-to-moderate changes in barcode abundance, possibly reflecting the suboptimal function of mammalian chaperones within yeast cells (Fig. 1e and Sup. Fig. 9a). Nevertheless, 20 significant interactions were identified, including rescue mediated by the primarily mitochondria-localized type III HSP40 chaperone, DNAJC11, and two membrane associated proteotoxicity models, TMEM106B and Kar2-beta-amyloid (Fig. 1f-g and Sup. Fig. 9b). These results are of interest as DNAJC11 mutant mice show prominent neuronal pathology with vacuolization of the endoplasmic reticulum and disruption to mitochondrial membranes^28^. Furthermore, both TMEM106b and beta-amyloid cell-based models have been associated with mitochondrial stress and dysfunction^23,29^. These findings suggest that the rescue with DNAJC11 may be related its ability to buffer against the effects of amylogenic proteins on mitochondrial function. Given the breadth of our screening it enabled the detection of a general trend of interaction between multiple human type II HSP40 chaperones (DNAJB1, DNAJB2, DNAJB4, DNAJB6, DNAJB8) and two yeast prions, RNQ1 and SUP35 within our library. These findings indicate that the misfolded intermediates produced by these two yeast prions may have similar properties, which is in agreement with data showing the ability of these two yeast prions to cross-seed each other’s aggregation^30^. Furthermore, the fact that SIS1, the yeast orthologue of these human DNAJB proteins, shows prominent activity against RNQ1 and SUP35, suggests that this class of misfolded prion species may be evolutionarily conserved substrates for the DNAJB family of chaperones^31^.

### Secondary validation of multiplexed screening results

To validate the interactions identified in the screen, we applied a testing paradigm similar to how the screen was conducted. For this purpose, we made use of a previously reported passaging-based growth assay, which we first verified against a set of known interactions (Sup. Fig 10)^32^. We then used the assay to individually validate two sets of hits, namely, those judged as statistically significant by our analytical pipeline and those with positive log2 fold changes that did not reach our significance threshold but represent “suspected interactions”. Hits identified as statistically significant with an FDR adjusted p-value of <0.05 were validated at a high rate, with 116/120 (96.7%) of these interactions reproducing upon individual testing (Sup. Table 3). We next verified a number of suspected interactions with positive log2 fold changes that did not meet statistical significance, with 59/95 (62.1%) of these suspected interactions showing rescue upon individual testing (Sup. Table 3). This indicates that the hit-calling algorithm can prioritize interactions that are likely to validate over using log2 fold changes alone, although some true hits may be missed as these likely do not survive our adjustments for multiple hypothesis testing. To estimate the sensitivity and specificity of the statistical pipeline, we assumed that the 175 validated interactions (116 significant and 59 suspected) represent most of the hits in the matrix of 5,850 interactions. Holding this to be the case, it suggests that our screening platform has an estimated sensitivity and specificity of ~66% and ~99%, respectively, which is on par or better than previously reported one-model-at-a-time screening approaches^33–36^. Furthermore, highlighting the power of our redundant barcoding strategy, if the number of barcodes analyzed for each model is progressively reduced from 5-7 redundant barcodes per model to 1 per model, a striking decrease in the number of hits captured is seen with each barcode removed (Sup. Fig. 11). These results reinforce the dramatic improvement in data quality afforded by transforming each model into multiple DNA-barcoded strains and analyzing their collective behavior.

To evaluate the performance of our platform against a larger, unbiased library of potential modifiers, we screened the hORFeome V8.1 human cDNA library against our pool. We chose to screen human ORFs in hopes of identifying hits that are more likely to directly translate to mammalian models of disease and because of a paucity of prior reports of human genes being tested in this paradigm, suggesting that many novel interactions likely remain to be uncovered. Screening ~900 members of the hORFeome library enabled the examination of ~35,000 interactions, representing ~9 times more interactions surveyed within our system than is typical^7,8,11,14,15^. Screening and subsequent validation of this collection of rescuers resulted in the identification of 54 confirmed genetic interactions (Sup. Table 4, Sup. Table 5). As expected with a library comprising random human genes, the occurrence of genetic interactions was significantly lower (0.14%) compared to the curated molecular chaperone screen (2.9%). Furthermore, when compared to other unbiased screens overexpressing yeast genes instead of human genes our hit rate remains lower (previously reported hit rates of 0.24%-1.13% vs. 0.14%)^8,10,11,14^. One of the main drivers of this difference is likely the failure of human genes to function in yeast cells, and further highlights the benefits of using a high-throughput quantitative screening approach to rapidly explore this sparsely populated interaction space^37^.

### DNAJB6 is a rescuer of multiple RNA-binding proteins implicated in ALS/FTD

During screening and subsequent validation, we identified a human chaperone, DNAJB6 that rescued the toxicity of cells expressing ALS/FTD-associated aggregation-prone RNA-binding proteins FUS, TDP-43, and hnRNPA1 (Fig. 2a). DNAJB6 is a type II HSP40 co-chaperone expressed in a number of tissues, including ubiquitously throughout the brain and spinal cord^38^. However, DNAJB6 expression is decreased in the brain with aging potentially sensitizing neurons to misfolded protein stress^39,40^. HSP40 co-chaperones comprise a large class of approximately 50 proteins in humans with an array of reported activities, but primarily function by binding to misfolded proteins, trafficking them to HSP70 chaperones, and regulating protein-protein interactions^41,42^. While DNAJB6 was the only type II HSP40 that was able to rescue TDP-43 toxicity, DNAJB1 and DNAJB2 could rescue the FUS and hnRNPA1 models, although their magnitude of rescue was 1/3 to 1/4 that of DNAJB6, suggesting DNAJB6 has properties that are unique among its family (Sup. Fig. 12). DNAJB6 has been previously shown to suppress the aggregation of polyglutamine repeat containing proteins^43–46^. In addition, mutations in DNAJB6 cause Limb Girdle Muscular Dystrophy D1 (LGMDD1), in which affected muscle tissue accumulates TDP-43 aggregates, suggesting it plays a role in the maintenance of TDP-43 solubility within humans^47,48^. Nonetheless, it remains unclear as to whether DNAJB6 is a broadly active modifier of other disease relevant clients and how DNAJB6 modulates misfolding^49,50^.

**Figure 2.**
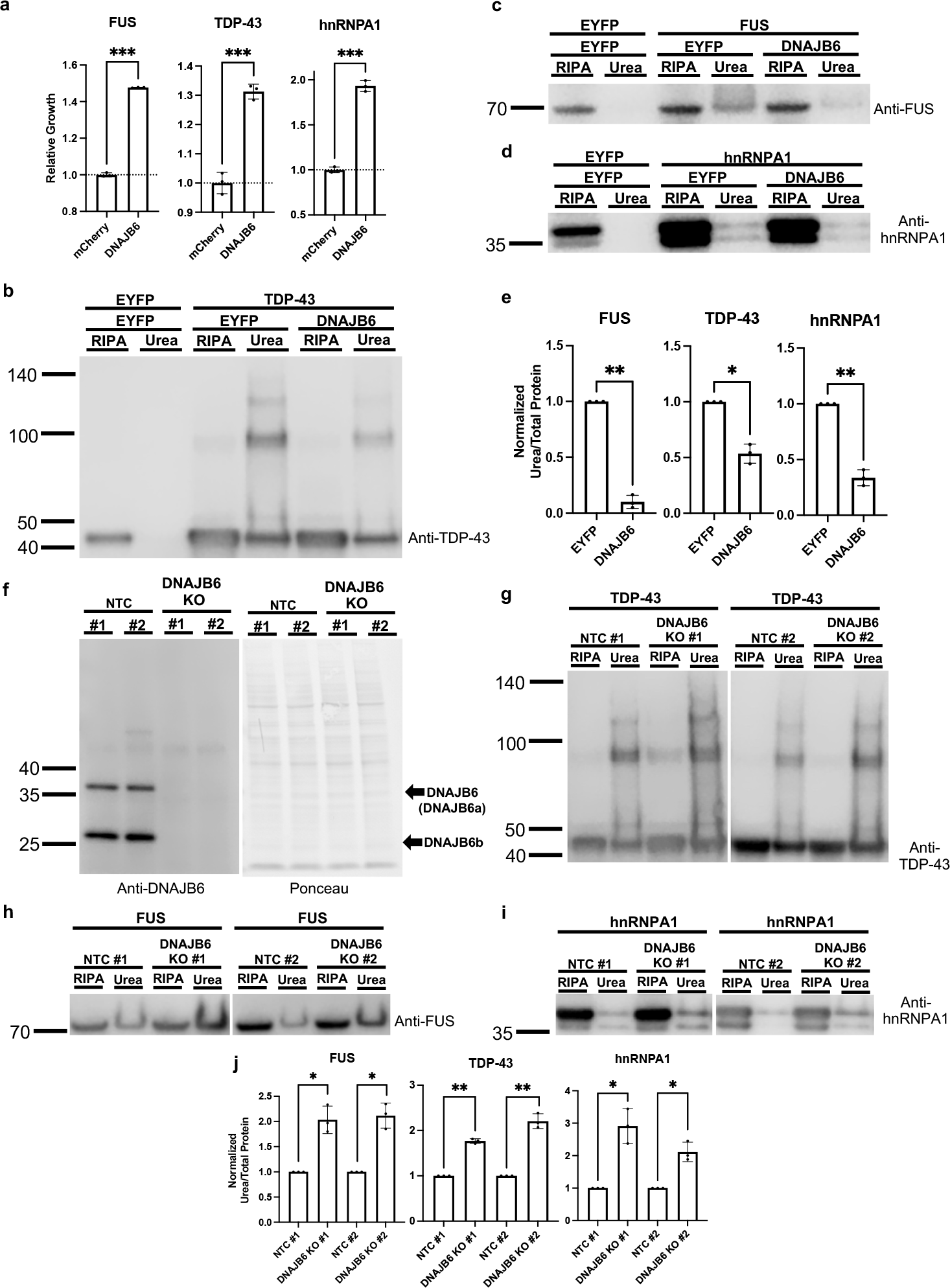
DNAJB6 is a rescuer of FUS, TDP-43, and hnRNPA1. **a**. DNAJB6 rescues proteotoxicity of FUS, TDP-43, and hnRNPA1 in yeast. Data are shown as mean ± s.d. for three biological replicates. **b-d**. Overexpression of FUS, TDP-43, and hnRNPA1 in HEK293T cells results in formation of SDS-insoluble (RIPA buffer insoluble), urea-soluble species.The formation of these species can be reduced by co-expression of DNAJB6. **e**. Quantification of the urea soluble species normalized to total protein detected by Ponceau S staining in b-d. Each independent replicate was normalized to its associated EYFP rescued sample. **f**. Validation of non-target control (NTC) and DNAJB6 KO lines. DNAJB6 has two isoforms; DNAJB6a and DNAJB6b, which are 36 kDa and 27 kDa in size, respectively. **g-i**. Overexpression of FUS, TDP-43, and hnRNPA1 in DNAJB6 KO lines shows an increase in the propensity to form SDS-insoluble, urea-soluble species. **j**. Quantification of the urea soluble species normalized to total protein detected by Ponceau S staining in g-j. Each independent replicate was normalized its associated EYFP rescued sample. All data are shown as mean ± s.d. for three biological replicates. All statistical tests were conducted with Welch’s t tests; ns = not significant P>0.05, *P≤0.05, **P≤0.01, ***P≤0.001, ****P≤0.0001.

In order to determine if the interaction between DNAJB6 and the RNA-binding proteins FUS, TDP-43, and hnRNPA1 are relevant within mammalian cell contexts, we employed an *in vivo* protein aggregation assay^49,51^. In this assay, when FUS, TDP-43, or hnRNPA1 are overexpressed within human embryonic kidney 293T (HEK293T) cells, they form SDS-insoluble species (RIPA buffer insoluble), that can be solubilized in urea. As compared to control cells co-transfected with enhanced yellow fluorescent protein (EYFP), cells co-transfected with DNAJB6 showed a reduction in the amount of SDS-insoluble FUS, TDP-43, and hnRNPA1 (Fig. 2b-e). This result demonstrates that DNAJB6 can reduce the formation of insoluble species for multiple RNA-binding proteins within mammalian cells.

### Endogenous DNAJB6 participates in the response to accumulation of insoluble FUS, TDP-43, and hnRNPA1

To further investigate the mechanisms and role DNAJB6 plays in the response to increasing cellular concentration of ALS/FTD-associated RNA-binding proteins, we transfected HEK293T cells with EYFP, FUS, or TDP-43 expression constructs and performed unbiased RNA-sequencing. As compared to the EYFP control condition, DNAJB6 was amongst the most strongly and significantly upregulated chaperones within the ~270 molecular chaperones observed upon overexpression of FUS and TDP-43 (Sup. Fig. 13).

To determine what role endogenous levels of DNAJB6 play in regulating FUS, TDP-43 and hnRNPA1 misfolding, Cas9 was used to generate multiple independent clones in which DNAJB6 was knocked out (Fig. 2f). In DNAJB6 knockout lines, SDS-insoluble FUS, TDP-43, or hnRNPA1 species were not observed when these proteins were expressed at endogenous levels (Sup. Fig. 14). However, upon overexpression of FUS, TDP-43, or hnRNPA1, greater amounts of SDS-insoluble species within DNAJB6 knockout lines were observed compared to the non-targeting gRNA control (NTC) lines (Fig. 2g-j).

Taken together, these data suggest that DNAJB6 is part of a programmed cellular response to rising levels of multiple aggregation-prone RNA-binding proteins and that physiological levels of DNAJB6 can regulate the solubility of these proteins.

### DNAJB6 undergoes phase separation and can prevent aberrant FUS interactions *in vitro*

The amino acid sequence of DNAJB6 contains multiple stretches of low complexity residues, which is a feature of many proteins that undergo liquid liquid phase separation (LLPS). This, together with the fact that DNAJB6 interacts with and rescues the aggregation of multiple proteins that undergo LLPS in cells (e.g. FUS, TDP-43), prompted us to investigate whether DNAJB6 might itself undergo LLPS^52–54^. To address this question, we deployed a previously published LLPS assay^51,55^. Incubation of DNAJB6 (3 μM or 0.25 μM) in near physiological salt concentrations (50 mM NaCl) resulted in phase separation and formation of liquid liquid droplets of DNAJB6 (Sup. Fig. 15a-b)^56^.

This experimental result raises the question as to whether DNAJB6 might co-partition with its phase separating client proteins such as FUS. To address this question, we mixed FUS and DNAJB6 at intracellular concentrations determined from the literature (1.5 μM and 0.25 μM respectively) in near physiological NaCl concentrations (50 mM)^57^. Under these experimental conditions, as well as at a 1:1 molecular ratio, (not shown) DNAJB6 and FUS co-partitioned into the condensed liquid droplet phase (Sup. Fig. 15c). Intriguingly however, the FUS + DNAJB6 condensates were more numerous, and had noticeably smaller diameters than FUS only condensates, suggesting that binding of DNAJB6 to FUS might modulate FUS-FUS interactions, thereby preventing FUS condensate progression and growth (Fig. 3a-b). If correct, this might also provide a mechanism through which DNAJB6 might modulate the propensity of FUS to undergo time-dependent progressive condensation (“aging”) into fibrillary aggregates^51,54,58^.

**Figure 3.**
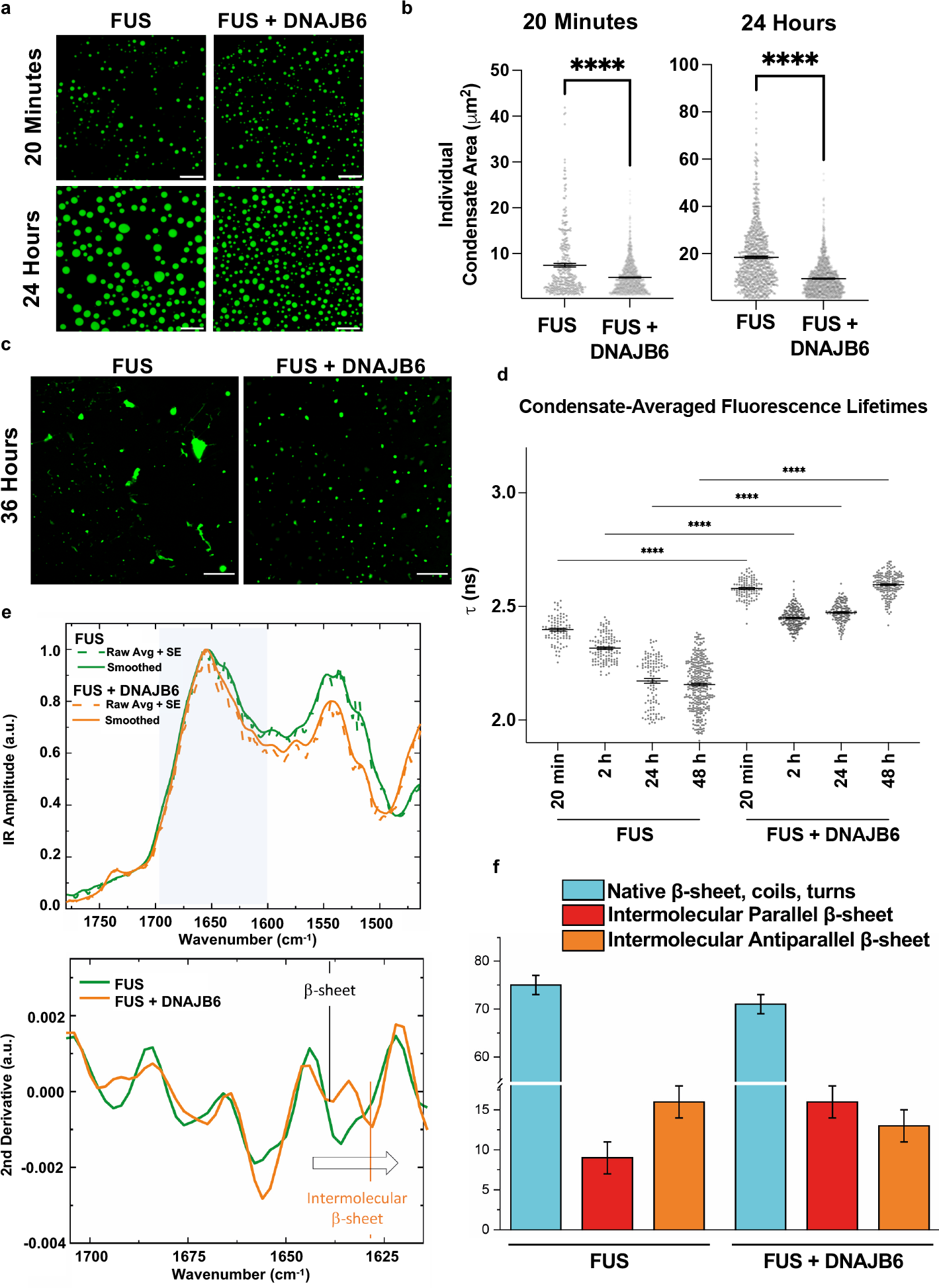
DNAJB6 modulates the dynamics and structure of FUS condensates. **a**. FUS and FUS + DNAJB6 condensates at 20 minutes and 24 hours. Scale bar = 10 μm. **b**. Quantification of condensate size as shown in panel a. 1 micron squared minimum size cutoff was used when quantifying condensates. Average condensate size was determined for multiple condensates within several different fields of view. At 20 minutes n=343 for FUS and n=1,455 for FUS + DNAJB6. At 24 hours, n=792 for FUS and n=1,459 for FUS + DNAJB6. Comparisons were conducted with Welch’s two-sided t test. ****P≤0.0001 Plots are shown as mean +/− SEM. **c**. FUS condensates form fibrillar aggregates when incubated for 36 hours. FUS condensates do not form fibrillar aggregates in the presence of DNAJB6 at 36 hours. Scale bar = 20 μm. **d**. Condensate-averaged fluorescence lifetimes. Results are based on 86 −326 condensates from 9 images taken over 3 independent experiments. One-way ANOVA (Holm-Sidak’s multiple comparison test), where **** is P<0.0001. **e**. IR average spectra from 3 independent FUS or FUS + DNAJB6 condensates and second derivative of the amide I band to deconvolve protein secondary structure contributions. **f**. Quantification of secondary structure within condensates by AFM-IR.

To further characterize the effect of DNAJB6 on FUS condensation across different timescales, we employed four orthogonal assays of the biophysical state of FUS – direct inspection of droplet morphology and fibrillary aggregates, fluorescence lifetime imaging microscopy (FLIM), fluorescence recovery after photobleaching (FRAP), and atomic force microscopy with infrared nanospectroscopy (AFM-IR). The direct inspection of the FUS condensates over time reveal that FUS + DNAJB6 condensates remain small and spherical, with no irregular fibrillary aggregates. In contrast, the FUS only condensates form large spherical condensates, which by 36 hours are accompanied by occasional fibrillary aggregates (Fig. 3c).

These results suggest that DNAJB6 constrains FUS condensation and prevents fibrillary aggregation. In addition, the persistently small size of the FUS + DNAJB6 condensates is consistent with the formation of a loose gel-like state which inhibits droplet droplet fusion events (Ostwald Ripening) that are characteristic of pure liquid liquid droplets. The FLIM assays, which measure the local packing environment showed a significant increase in FLIM lifetimes of FUS within FUS + DNAJB6 condensates at all time points from 20 minutes to 48 hours after condensate formation compared to the FUS only control (Fig. 3d). This result suggests that DNAJB6 permits FUS condensation to occur, but limits FUS-FUS packing and further condensation. In good agreement with this conclusion, the FRAP experiments at 30 minutes, revealed increased FRAP recovery rates in FUS + DNAJB6 condensates (Sup. Fig. 15d). However, at 24 hours, both the larger FUS only condensates and the smaller FUS + DNAJB6 condensates showed little FRAP recovery (data not shown). This likely reflects further condensation of FUS in both condensate types, albeit with potentially different condensation states in the FUS only versus the FUS + DNAJB6 condensates at these later time points that are not discriminable by FRAP.

The morphological analysis of condensate size, together with the FLIM and FRAP data suggest that the FUS + DNAJB6 condensates may form spherical, loosely condensed gel-like structures. In contrast, FUS-only condensates tended to form more compact condensates with a propensity to form fibrillar aggregates.

To interrogate the nature of the spherical condensates in the FUS-only and the FUS + DNAJB6 conditions, we applied atomic force nano infrared spectroscopy as described previously^51,55^. Studies of individual spherical condensates on zinc selenide chips revealed that spherical FUS + DNAJB6 condensates contained higher content of intermolecular parallel β-sheet and less content of intermolecular antiparallel β-sheet than the spherical FUS-only droplets (Fig. 3e-f and Sup. Fig 16). Furthermore, the parallel β-sheet of FUS + DNAJB6 shifted at a lower wavenumber, indicating more extended strands and more hydrogen bonding. These spectral properties have previously been associated with gelled polymers and therefore support the hypothesis that DNAJB6 prevents progression of FUS condensation into irreversible fibrillary aggregates by incorporating them into looser gel-like condensates^55,59–64^.

### Characterization of DNAJB6 via domain deletions and deep mutational scanning

Having established the ability of DNAJB6 to phase separate and directly interact with its clients to prevent their misfolding, we sought to decipher the regions within the protein required for its effect within cells. A series of DNAJB6 deletion mutants were constructed, including versions lacking the J-domain which is necessary for activation of HSP70 partners, the glycine/phenylalanine rich region which contributes to its ability to phase separate, and the serine rich domain that is implicated in client recognition and binding^43,45^. Deletion of any of these domains prevented DNAJB6 from rescuing FUS toxicity within yeast, consistent with the recent results using purified proteins for a related HSP40 chaperone, DNAJB1, and its interaction with FUS (Sup. Fig. 17a)^65^. All the examined deletion mutants were expressed in yeast except for the variant with the J-domain removed. To better examine the role for the J-domain a point mutant within the conserved HPD motif of the J-domain (H31Q) was created. This mutant expresses well but blocks the ability of DNAJB6 to stimulate the ATPase activity of HSP70 family members^42^. The H31Q mutation also rendered DNAJB6 non-functional for rescuing FUS-mediated toxicity. These results suggest that all domains within DNAJB6 are required for its activity and that cooperating with a HSP70 partner may be necessary for its full function *in vivo* (Sup. Fig. 17a-b).

During our deletion analysis, we observed that loss of the glycine/phenylalanine (G/F) rich domain or the serine (S) rich domain from DNAJB6 enhanced the toxicity of the FUS model (Sup. Fig. 17b). These regions have been implicated in substrate recognition and are also the site of mutation within LGMDD1 patients along with multiple variants of uncertain significance within the ClinVar database. To decipher the function of this critical region at increased resolution, we conducted a deep mutational scan (DMS) across the 112 amino acids of the G/F and S-rich regions. Deep mutational scanning libraries were constructed by performing comprehensive mutagenesis of each codon, yielding versions of DNAJB6 with all possible amino acids or a stop codon at each interrogated position (Fig. 4a). The library of variants was then tested for their ability to rescue the proteotoxicity induced upon FUS overexpression, creating a comprehensive fitness landscape of all single amino acid mutations in the G/F and S rich region on DNAJB6 activity (Fig. 4b). Mutagenesis libraries were constructed and screened in biological duplicates, with strong correlation between replicates observed (see Materials and Methods, Sup. Fig. 18). Notably, regions rich in patient mutations causing LGMDD1 (amino acids 89-100) frequently resulted in a reduction in activity, with subsequent validation of these mutations confirming a loss of activity compared to wild-type (WT) DNAJB6 against the FUS model (Fig. 4b-c). Similar results were obtained when these mutants were tested against the TDP-43 model, in agreement with the existence of a common mechanism of interaction between DNAJB6 and its aggregation-prone clients (Fig. 4c). Numerous variants of uncertain significance (VUS) in DNAJB6 present in ClinVar were also captured within the data. Follow-up studies individually examining these mutants revealed a subset which were defective in their ability to rescue FUS and TDP-43 expressing models, suggesting that additional disease associated mutants may exist outside what is currently annotated (Fig. 4c). Within the DNAJB6 mutational landscape, variants with enhanced activity against FUS were observed. Outside of the conservative S192T mutation which showed one of the strongest effects on rescue, a general trend was seen where mutation to an acidic residue in stretches between 138-171 and 182-189 appeared to enhance activity compared to WT DNAJB6 (Fig. 4b). In line with this observation, the single acidic amino acid within these stretches, D158, was critical for function as mutation of D158 to almost any other amino acid other than glutamic acid reduced the activity of the protein (Fig. 4b). Based on the screening results, several DNAJB6 mutants with enhanced activity were selected for validation against both the FUS and TDP-43 models within yeast, revealing 3 mutations that showed clear gains in activity as compared to the wild-type protein (Fig. 4c).

**Figure 4.**
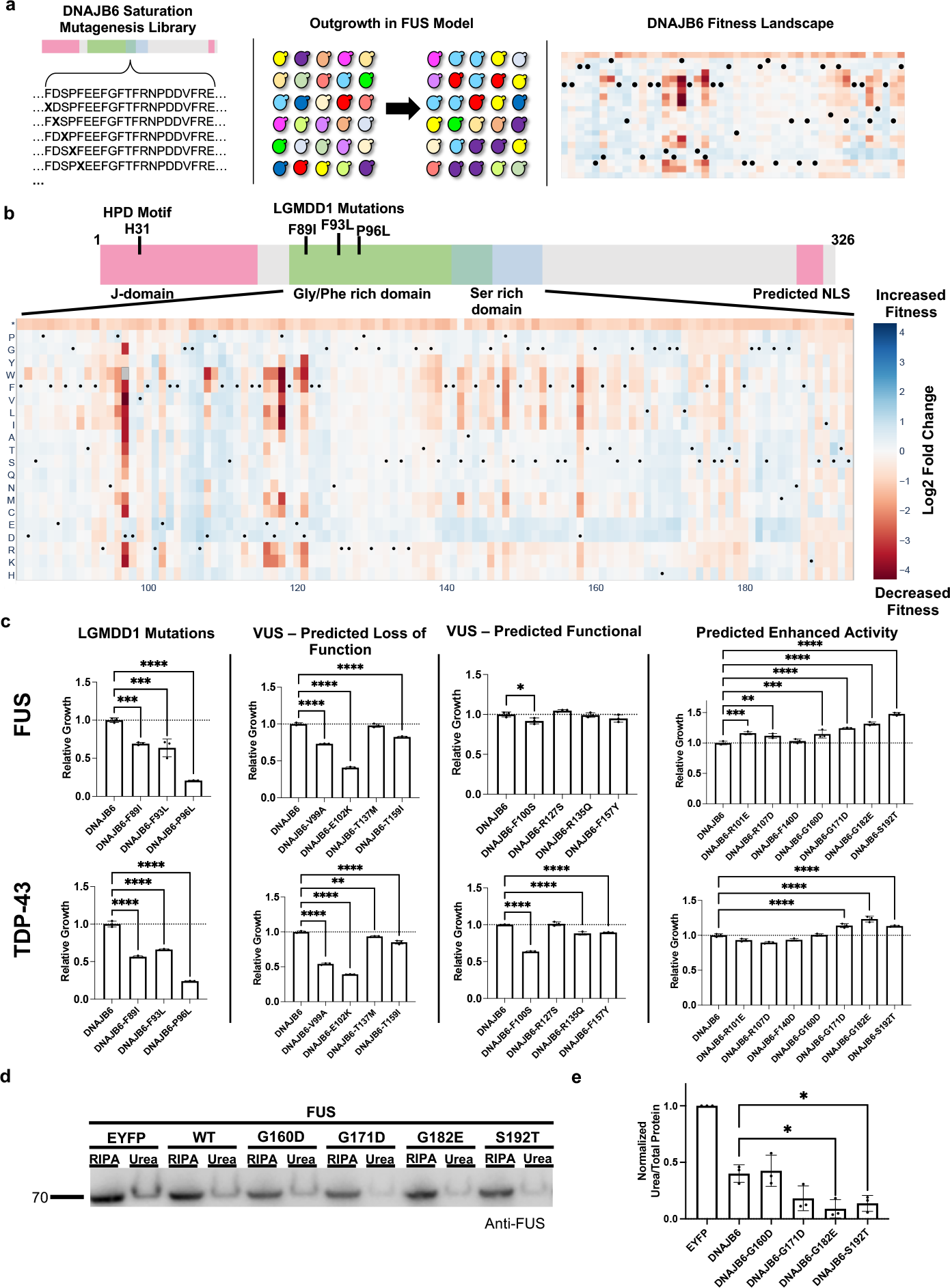
Deep Mutational Scan (DMS) of DNAJB6 identifies potentiated variants. **a**. Overview of DMS approach for testing DNAJB6 variants against the FUS model. A library of DNAJB6 mutants is generated using a site directed mutagenesis-based approach, the resulting plasmid library is transformed into yeast containing the FUS model and grown under inducing conditions, finally the relative growth rates of all mutants are determined and normalized to the wild-type DNAJB6 variant. **b**. DMS heatmap for residues 83-194 of DNAJB6. Intensity of blue or red colored boxes indicates increased or decreased activity as compared to the wild-type DNAJB6 protein. * indicates stop codon mutation, black dots (•) mark the wild-type residue for each site in the protein. **c**. Validation of DMS results in the FUS model and testing of variants against the TDP-43 model. **d**. Testing of potentiated DNAJB6 variants in mammalian cells for their ability to reduce SDS-insoluble, urea-soluble FUS species upon overexpression. **e**. Quantification of the urea soluble species normalized to total protein detected by Ponceau S staining in d. Each independent replicate was normalized its associated EYFP rescued sample. All data are shown as mean ± s.d. for three biological replicates. Comparisons were conducted with ordinary one-way ANOVA with display of significant comparisons; *P≤0.05, **P≤0.01, ***P≤0.001, ****P≤0.0001.

To further establish the relevance of the enhanced DNAJB6 variants, their ability to reduce the formation of SDS-insoluble, urea-soluble FUS species in mammalian HEK293T cells was examined. Given that WT DNAJB6 almost entirely reduced the formation of SDS-insoluble, urea-soluble FUS, the assay was modified by transfecting more FUS expression plasmid into cells which resulted in higher levels of SDS-insoluble, urea-soluble FUS species (Sup. Fig. 19). Utilizing higher FUS expression, the potentiated DNAJB6 variants, G182E and S192T showed a further reduction in the formation of SDS-insoluble, urea-soluble FUS species as compared to the wild-type protein (Fig. 4d-e). These findings demonstrate that the activity of DNAJB6 can be further improved and that rescuer proteins optimized in yeast can be readily translated to mammalian cell systems^27^.

## Discussion

Modeling of neurodegeneration in simplified cellular systems has yielded insights into how proteotoxic aggregation-prone proteins disrupt cellular function. However, a great deal of additional information is required to fully understand these complex processes, and provide practical information that can be translated into effective diagnostics and therapeutics for the associated human medical disorders. The work described here advances this goal in two ways. First, we developed a multiplexed approach in which numerous misfolding-prone proteins can be screened in parallel to identify a rich dataset of candidate modulators acting either in a protein-specific way, or in a more general class-specific way. Second, our work has uncovered a previously unrecognized mechanism whereby protein:protein interactions can modulate the propensity of misfolding-prone proteins to form species that injure cells – namely by the formation of gel-like biomolecular condensates instead of irreversible fibrillar condensates. In the paragraphs below, we explore each of these concepts in greater detail.

### A novel platform for identifying modulators of proteotoxic protein misfolding

Our platform provides several advantages over conventional approaches. Specifically, by inserting the same misfolding-prone protein into several uniquely barcoded strains and by analyzing their collective response to a given genetic perturbation, we are able to greatly enhance our assay sensitivity and specificity. Crucially, the quantitative nature of our approach better enables us to capture mild changes in growth by a putative rescuer as compared to traditional semi-quantitative methods. Moreover, by simultaneously studying multiple models within the same genetic background and under the same testing paradigm, we are able to make broad observations about the nature of rescuers and the relationship between models. Finally, in contrast to previous studies that screen cDNA libraries derived from the model organism being used (e.g. yeast or flies), we demonstrate across dozens of models that our platform enables direct screening of human genes within simple organisms such as yeast to identify disease relevant suppressors. Overall, this work establishes a high-throughput platform for identifying novel genetic suppressors applicable to any family of proteins so long as their expression in yeast causes a growth defect that is dependent upon the biological function the user desires to interrogate^66,67^. Furthermore, with minor modifications our approach can be readily applied toward screening against libraries of small molecules to identify compounds with specific versus broad activities against classes of disease relevant proteins such as the NDD-associated proteins studied in this work.

An additional finding from our screens was the lack of concordance between proteotoxic species of the same class, such as various dipeptide repeats and poly-alanine models. In contrast, RNA-binding proteins showed a general trend for having rescuers that were active on several members of its group. No rescuers, however, showed clear activity on all RNA-binding proteins and each tended to show a preference for a particular subset (Fig. 2 and Sup. Fig. 8-9). These results imply that there is no universal feature that is conserved across a given family of NDD-associated proteins, but that there can still be some areas of conserved identity among the misfolded species that enable genetic modifiers to interact with several members of the family.

As our library also contained variants of the same protein of interest with different patient mutations, we asked what, if any, was the effect of these mutations on our screening results. Interestingly, despite many of the tested mutations having been shown to accelerate the rate of protein misfolding or drastically alter the localization of the variant protein, they appeared to be rescued in a similar manner as the wild-type protein (Sup. Fig. 8-9)^51,54,68,69^. These results suggest that the tested patient mutations, while increasing the probability of misfolding (and thus probability of disease), do not appear to fundamentally change the underlying mechanism of toxicity or the misfolded state that is present within the cell. These results raise the possibility that a therapeutic modality designed against the wild-type form of a protein might also work for patients with rare disease-causing variants.

### Changing the phase state of phase separating proteins forming biomolecular condensates

The work reported here has revealed that certain chaperone proteins (e.g. DNAJB6) can interact with phase separating RNA binding proteins (e.g. FUS, TDP-43, hnRNPA1). Prior work has shown that these RNA binding proteins form reversible biomolecular condensates that are crucial to their function in nuclear RNA transcription, and particularly in cytoplasmic RNA translation in selected vulnerable niches such as axon terminals of neurons^51,55,70–72^. However, these biomolecular condensates, even when comprised of wild-type RNA binding proteins, have a propensity to condense into irreversible assemblies enriched in β-sheet fibrils^51,54,58^. The formation of these irreversibly condensed assemblies inhibits the function of the RNA binding proteins in RNA transcription and RNA translation. This latter effect reduces new protein synthesis in axon terminals, which are particularly dependent upon local RNA translation for the production of niche-specific proteins involved in synaptic viability and function^51,55,70–72^.

Our data shows that DNAJB6 co-fractionates with FUS under near physiological conditions, and encourages the formation of less compact, gel-like condensates, thereby preventing the formation of irreversible fibrillar aggregated species. This finding is in contrast to other identified rescuers of FUS misfolding, such as TNPO1, which deters FUS aggregation by dissolving condensates^55^.

Additional work will now be required to probe if and how chaperone-induced formation of gel-like condensates might still allow effective local RNA translation in neurons. For instance, do the chaperone-induced gel-like condensates act like a percolated gel, and permit release of naked RNA species for access by local RNA translation machinery? Alternatively, are the chaperone-induced gel-like condensates simply being held in a phase state that is still reversible, perhaps through interactions with other local chaperones and disaggregases, converting them back to more liquid-like assemblies? In this latter regard, while our work has focused primarily on DNAJB6, HSP40 family members often work in concert with HSP70 chaperones, which are also known to regulate the solubility of ALS/FTD-associated RNA-binding proteins^73^. In future studies, modeling *in vitro* the behavior of DNAJB6 with a HSP70 partner could reveal additional insights into the process by which it helps maintain aggregation-prone proteins such as FUS soluble *in vivo*.

Finally, it is important not to overlook prior work which has demonstrated that DNAJB6 can prevent the aggregation of poly-glutamine repeat expansion containing proteins, along with acting on beta-amyloid and alpha-synuclein^39,43,44,46,74,75^. Taken together with our study, these results raise the possibility that DNAJB6 represents a generalized chaperone for a number of clinically relevant misfolded species. Furthermore, the ability to observe DNAJB6 activity in an easy to manipulate yeast model enables the high-throughput optimization of its activity. We demonstrate that extensive deep mutational scanning of DNAJB6 can identify enhanced variants. Additional screening of these mutant libraries against numerous protein misfolding models could be used to select for more broadly-active versions of the protein. Alternatively, these same mutant libraries could be used to selectively tune DNAJB6 towards a narrower class of targets should an increase in specificity be desired for eventual therapeutic applications. By examining, the properties and three-dimensional structure of the obtained enhanced DNAJB6 proteins, we can gain fundamental insights into how these classes of proteins may function and evolve, along with informing future gene based therapies or the design of small molecules that activate DNAJB6 function or mimic its activity.

## Materials and Methods

### Yeast Strains and Media

Barcoded *S. cerevisiae* yeast BY4741 *MATa his3*Δ*1 leu2*Δ*0 ura3*Δ*0 met15*Δ*0* strains were purchased from Horizon (Cat. #YSC5117). To introduce rescuers through mating, rescuer containing *S. cerevisiae* yeast BY4742 strains *MATɑ his3*Δ*1 leu2*Δ*0 ura3*Δ*0 lys2*Δ*0* were used. Individual barcoded strains containing expression vectors were maintained in Synthetic Complete (SC) -ura media (20 g/L glucose, 1.5 g/L Drop Out mix [US Biological D0539-09A], 1.7 g/L Yeast Nitrogen Base [US Biological Y2030], 5 g/L Ammonium Sulfate [Fisher H8N2O45], supplemented with 18 mg/L Leucine and 9 mg/L Histidine). Individual rescuer BY4742 strains were maintained in SC -his media (20 g/L glucose, 1.5 g/L Drop Out mix [US Biological D0539-09A], 1.7 g/L Yeast Nitrogen Base [US Biological Y2030], 5 g/L Ammonium Sulfate [Fisher H8N2O45], supplemented with 18 mg/L Leucine and 1.8 mg/L Uracil). Mating was conducted in YPD media (20 g/L glucose, 20 g/L peptone, and 10 g/L yeast extract). Selection for mated strains was conducted in SC -ura -his media (20 g/L glucose, 1.5 g/L Drop Out mix [US Biological D0539-09A], 1.7 g/L Yeast Nitrogen Base [US Biological Y2030], 5 g/L Ammonium Sulfate [Fisher H8N2O45], supplemented with 18 mg/L Leucine), while outgrowth of induced mated strains was carried out in SC -ura -his gal media (20 g/L galactose, 1.5 g/L Drop Out mix [US Biological D0539-09A], 1.7 g/L Yeast Nitrogen Base [US Biological Y2030], 5 g/L Ammonium Sulfate [Fisher H8N2O45], supplemented with 18 mg/L Leucine)

### Plasmids

Proteotoxic genes and controls were cloned into either the pAG416GAL-ccdb (Addgene #14147) or pAG426GAL-ccdb (Addgene #14155) using Gateway LR II Clonase Enzyme mix (Invitrogen). Once expression plasmids were sequence verified, they were transformed into barcoded BY4741 strains using standard LiOAc transformation protocols and plated on SC -ura agar plates. Yeast rescuer genes and control rescuer genes were cloned into pAG413GAL-ccdb (addgene #14141) using Gateway cloning. Human rescuer genes from the hOrfeome V8.1 Library collection were cloned into a derivative of pAG413GAL-ccdb, pAG413GAL-ccdb-6Stop, wherein the 3’ attR2 site was modified to encode a stop codon 6 amino acids downstream of the last codon to compensate for a lack of a stop codon in the ORFeome.

All mammalian expression vectors were cloned into the pLEX307 backbone (Addgene #41392) using Gateway LR II Clonase Enzyme mix (Invitrogen).

Plasmid DNA was isolated using standard miniprep buffers (Omega Biotek) and silica membrane columns (Biobasic). All expression plasmids were Sanger sequenced to confirm the appropriate insert (Genewiz).

### Yeast Multiplexed Screening

Each plate of rescuers was screened in biological duplicates. A fresh aliquot (500 μL) of frozen barcoded yeast pool was inoculated into 5 mL of SC -ura media and rotated at 30°C. At the same time, 5 μL of each rescuer strain was inoculated into 500 μL of SC -his media in 96 well 2 mL deep well plate format (VWR) and shaken at 900 rpm at 30°C. 24 h later, 5 μL of the saturated barcoded yeast pool was mixed individually with 5 μL of rescuer strain in a new 96 well plate where each well was filled with 500 μL of YPD and shaken at 900 rpm at 30°C. For selection of mated strains, 20 h later, 5 μL of mated barcoded yeast pool was transferred into a new 2 mL deep well plate filled with 500 μL of SC -ura -his media and shaken at 900 rpm at 30°C for 24 h. For outgrowth, 2 μL of the mated and selected pool was inoculated into 1 mL of SC -ura -his galactose media and shaken at 1,000 rpm at 30°C for 30 h.

After growth, 100 μL of yeast culture was removed and the optical density (OD595) of the culture was determined in a 96 well plate reader (Tecan). After measurement of culture density, genomic DNA was extracted using a modified LiOAc-SDS extraction method. Briefly, plates were centrifuged for 5 min at 4,000 rpm. Supernatant was discarded and the pellet was resuspended in 200 μL of 200 mM LiOAc with 1% SDS with rigorous pipetting. Plates were sealed with aluminum foil and incubated at 70°C for 20 min to enable lysis. 600 μL 100% ethanol was added to each well and pipetted up and down rigorously before being centrifuged for 10 min at 4,000 rpm. Supernatant was discarded and pellets were air dried for 30 min under flame. Pellets were then resuspended in 200 μL 1X TE and incubated at 42°C for 30 min. The plates were centrifuged for 10 min at 4,000 rpm and the supernatant containing DNA was pipetted into a new plate for storage at −20°C. Raw sequencing reads from the chaperone screen have been uploaded to the NCBI SRA under BioProject PRJNA769721 (SUB10508463). Raw sequencing reads from the orfeome screen have been uploaded to the NCBI SRA under BioProject PRJNA769721 (SUB10508562).

### Sequencing Library Preparation

For sequencing on NextSeq 500/550 (Illumina), libraries were prepared from genomic DNA in two PCR steps. The first step amplifies genomic DNA containing the DNA barcode and attaches an internal index to designate which column the well was amplified from. The second PCR attaches Illumina indexes to the amplicon, wherein the combination of Illumina indexes indicates the row and plate location of the well. The first PCR step was done in technical duplicates unless otherwise stated with Taq polymerase (Enzymatics). The following reaction mix was used: 2 μL 10X Taq buffer, 0.1 μL 100 μM forward primer, 0.1 μL 100 μM reverse primer, 0.1 μL Taq polymerase, 0.4 μL 10 mM dNTPs, 0.5 μL DNA, and 16.8 μL H_2_O. The following cycling conditions were used: 1. 94°C, 180 s, 2. 94°C, 30 s, 3. 60°C, 20 s, 4. 72°C, 30 s, 5. Return to step 2 27X, 6. 72°C, 180 s. After the first round of PCR, technical replicates of each individual well were pooled. For the second round of PCR where Illumina indexes were attached, the following reaction mix was used: 2 μL 10X Taq buffer, 0.1 μL 100 μM forward primer, 0.1 μL 100 μM reverse primer, 0.1 μL Taq polymerase, 0.4 μL 10 mM dNTPs, 0.5 μL DNA from first round PCR, and 16.8 μL H2O. The following cycling conditions were used: 1. 94°C, 180 s, 2. 94°C, 30 s, 3. 56°C, 20 s, 4. 72°C, 30 s, 5. Return to step 2 7X, 6. 72°C, 180 s. After the second round PCR, all reactions corresponding to a plate of screening were pooled together. The reaction products were run out on a gel and a band corresponding to the right size was gel extracted. Libraries were quantified with the NEBNext Library Quant Kit for Illumina according to manufacturer instructions (NEB). Pooled libraries were combined and sequenced with a 75 cycles NextSeq 500/550 High Output Kit on a NextSeq 500/550 machine (Illumina).

### Analysis of Multiplexed Screening

Raw reads in fastq format were trimmed and assigned to wells via combinations of Illumina indexes and column designating internal indexes. 20 bp barcode sequences were aligned to a reference genome allowing for +1 or −1 shifts in the sequencing phase using bowtie2. Raw counts of exact matches for each barcode were determined and well-read counts were normalized by the total number of reads in that well and converted to counts per million (CPM) unless otherwise stated. Wells were analyzed in batches with other wells in the same plate. Wells with less than 15,000 total reads were discarded in addition to wells where 1 biological replicated received less than 15,000 total reads. After CPM normalization, the estimated actual abundance of reads were calculated by normalizing against the optical density (OD595) of that well, which was measured immediately prior to harvesting with a 96 well Infinite F50 plate reader (Tecan). Wells containing control or inert rescuers were identified and the average read counts of each barcode in controls was determined. The variance in the number of reads between control wells for each barcode was determined. The mean-variance relationship was modeled using the equation log(σvariance-mean) = log(k)+b*log(mean)σ as previously described^76^. The barcode mean and adjusted variance were used to determine whether a barcode in a test well was significantly upregulated using a one-sided cumulative density function assuming a normal distribution. The associated p-value was adjusted using a Benjamini-Hochberg procedure correcting for the number of tests in that well. To obtain model level information from individual barcode strains, p-values from independent barcode strains associated with the same model were combined using Stouffer’s method. After this summary value was obtained, a further Benjamini-Hochberg procedure was used to correct summary values for the number of models and the number of wells in each plate. Average log2 fold change was calculated as the base 2 logarithm of the average change in counts over the expected value in the control wells. Analysis was conducted with custom scripts in R Version 4.0.2.

### Spot Assays

Yeast strains to be assayed were grown overnight in selective synthetic complete media until saturation was reached. Saturated cultures were serially diluted 1:5 in sterile PBS (Gibco). To SC –ura -his glucose or galactose agar plates, 5 μL of diluted culture was spotted. Plates were left for 30 minutes to dry before being inverted and incubated at 30°C for 48 h, followed by being scanned to document growth.

### Yeast Liquid Culture Growth Assay

Proteotoxic yeast strains were grown in 500 μL SC -ura media in plate format shaken at 1,000 rpm at 30°C for 24 h. At the same time, rescuer yeast strains were grown in 500 μL SC -his media shaken at 1,000 rpm at 30°C for 24 h. After growth, 5 μL of appropriate proteotoxic and rescuer yeast strains were mixed in 500 μL YPD and shaken at 1,000 rpm at 30°C for 24 h. To 500 μL of dual selective SC -ura -his media, 5 μL of mated strains were inoculated and shaken at 1,000 rpm at 30°C for 30 h. 48 h prior to reading, mated and selected strains were inoculated in SC -ura -his galactose media at one of 3 dilution factors (Sup. Table 1) depending on their growth rate and shaken at 1,000 rpm at 30°C for 24 h. 24 h after initial inoculation, strains were passaged depending on their specified dilution factor into fresh SC -ura -his galactose media. Upon reaching the assay endpoint, 100 μL of each well was transferred to a 96 well plate (Greiner) and the optical density was determined on a 96 well plate reader (Tecan). Multiple media only wells were also quantified as a baseline and these values were subtracted from optical density measurements. All statistics were performed in GraphPad Prism Version 9.2.0.

### Protein Harvesting and Western Blotting

#### RIPA/Urea Extractions

Cells in 24-well dishes were washed with ice cold PBS (Gibco) after media was removed. 250 μL of RIPA buffer (50mM Tris-HCl (pH 7.4), 150mM NaCl 1% NP-40, 0.5% sodium deoxycholate and 0.1% SDS, Alfa Aesar) was added to each well and allowed to sit for 2 min. Cells were resuspended in RIPA buffer and moved to conical tubes. To lyse cells further, cells were sonicated for 10 s while kept on ice. Cell lysate was centrifuged for 20 min at 12,000 × g at 4°C and supernatant was saved as RIPA soluble fraction. Pellets were washed 1X with RIPA buffer and 50 μL urea buffer was added (8M Urea, 2M Thiourea, 4% CHAPS,) with vigorous pipetting to resuspend pellet and spun for 20 min at 12,000 × g at 4°C. The concentration of protein in the RIPA buffer was determined with a Bradford assay and lysates were adjusted to a final concentration between 250-350 μg/μL depending on the yield of the lowest concentration of the lysate in the set in which it was processed with 1X LDS loading buffer (Invitrogen). RIPA soluble fractions were boiled for 5 minutes and stored at −80°C until use. Urea fractions were stored at −80°C without boiling.

#### Yeast Lysate Extraction

Cells were collected and centrifuged at 2,300 × g for 2 min and washed with 1 mL of dH_2_O. To pelleted yeast cells, 200 μL of 0.1M NaOH was added and cells were resuspended by vortexing. Cells were allowed to lyse for 10 min at room temperature. Lysed cells were spun at 13,000 × g for 1 min and supernatant was discarded. Pellets were resuspended in 50 μL of dH_2_O and 25 μL of 200 mM DTT (Fisher) was added along with 25 μL of 4× LDS loading buffer (Invitrogen). Samples were boiled at 95°C for 5 min and subsequently centrifuged at 800 x g for 10 min at 4°C. Supernatants were collected and moved to a new tube for storage at −20°C.

#### Western Blotting

10 μL of normalized lysate with loading buffer was loaded into NuPAGE 4 to 12% Bis-Tris protein gels (Invitrogen) and subjected to 100 V electrophoresis for 65 min. Separated proteins were transferred onto a 0.2 μM PVDF membrane and blocked with SuperBlock (Invitrogen). Primary antibodies were diluted in SuperBlock with 0.1% Tween-20 and incubated overnight at 4°C with gentle rotation. Blots were washed with TBST before secondary antibody incubation. Blots were imaged with the Odyssey XF imaging system (Li-Cor) using the chemiluminescent detection. After transfer, blots were stained with Ponceau S stain (G Biosciences) for 15 min. Total protein was imaged on a LAS-4000 imager (Fujifilm). Band intensities were quantified with Image Studio Lite (Li-Cor).

TDP-43 was detected with a polyclonal rabbit antibody at a 1:2,500 dilution (Proteintech 10782-2-AP). FUS was detected with a polyclonal rabbit antibody at a 1:2,500 dilution (Proteintech 11570-1-AP). hnRNPA1 was detected with a polyclonal rabbit antibody at a 1:5,000 dilution (Proteintech 11176-1-AP). DNAJB6 was detected with a monoclonal mouse antibody at a 1:2,500 dilution (Proteintech 66587-1-Ig). A goat anti-rabbit HRP conjugated antibody was used at a 1:50,000 dilution (Invitrogen G21234). A goat anti-mouse HRP conjugated antibody was used at a 1:10,000 dilution (Invitrogen 31430). All statistics were performed in GraphPad Prism Version 9.2.0.

### Protein purification and *in vitro* LLPS experiments

FUS-mEmerald was purified as previously described^55^. His-Sumo tagged DNAJB6 expression vector was transformed into *E. coli* BL21(DE3) (NEB) for protein expression and purification. DNAJB6 expressing cells were grown at 37°C until an OD600 of 0.6 was reached. Expression was induced with 0.5 mM IPTG overnight at 16°C. Cell pellets were collected and subjected to high pressure lysis (Constant System) in lysis buffer (50 mMTris pH 7.5, 20 mM imidazole, 500 mM NaCl with 1× protease inhibitor cocktail). Lysate was centrifuged at 100,000 × g and collected supernatant was applied to a 10 mL Ni-Advance (BioServ, UK) column. After washing, His-Sumo tagged DNAJB6 was eluted in 250 mM imidazole containing buffer and cleaved overnight with ULP-protease at 4°C. Cleaved DNAJB6 was diluted 5-fold before running through a cation exchange column. SP Sepharose chromatography was conducted in 50 mM HEPES, pH 7.5 with a salt gradient from 5 M – 1 M NaCl. Fractions containing the protein were concentrated and subjected to size-exclusion on a Superdex-75 16/600 column in 50 mM HEPES and 100 mM NaCl, pH 7.5. Through all stages of purification, presence of DNAJB6 was monitored via SDS-PAGE.

DNAJB6 was labeled with Alexa Fluor™ 555 C2 Maleimide (Thermo Scientific) following the manufacturer’s guidelines.

For LLPS experiments, concentrated, purified proteins (FUS, DNAJB6, and/or BSA) were diluted to 1.5 μM in a 50 mM NaCl solution with 50mM Tris pH7.5 unless otherwise stated. For imaging, condensates were maintained in PEG-silane coated Ibidi™ coverslides to avoid wetting. Imaging was conducted with Zeiss Axiovert 200M microscope with Improvision Openlab software using 100X magnification objective.

### TCSPC-FLIM

mEmerald-tagged WT-FUS in NaCl solution (and DNAJB6) were mixed in milli-Q water, to give final protein concentrations of 1 μM of WT-FUS, 0.16 μM WT-DNAJB6, and 60 mM NaCl. 7 μL of each condensate mixture was deposited in individual silicon wells (Press-to-Seal, ThermoFisher Scientific) attached on 1.5 thickness coverslips (Superior Marienfeld, Lauda-Konigshofen, Germany) for ageing and imaging. Samples were imaged on a home-built confocal fluorescence microscope equipped with a time-correlated single photon counting (TCSPC) module. A pulsed, supercontinuum laser (Fianium Whitelase, NKT Photonics, Copenhagen, Denmark) provided excitation a repetition rate of 40 MHz. This was passed into a commercial microscope frame (IX83, Olympus, Tokyo, Japan) through a 60× oil objective (PlanApo 60XOSC2, 1.4 NA, Olympus). The excitation and emission beams are filtered through GFP-appropriate bandpass filters centered at 474 and 542 (FF01-474/27-25, FF01-542/27, Semrock Inc., NY, USA). Laser scanning was performed using a galvanometric mirror system (Quadscanner, Aberrior, Gottingen, Germany). Emission photons were collected on a photon multiplier tube (PMT, PMC150, B&H GmBH, Berlin, Germany) and relayed to a time-correlated single photon counting card (SPC830, B&H GmBH). Images were acquired at 256×256 pixels for 120 s (i.e., 10 cycles of 12 s). Photon counts were kept below 1% of laser emission photon (i.e., SYNC) rates to prevent photon pile-up. TCSPC images were analysed using an in-house, MATLAB-based (MathWorks, Natnick, MA, USA) phasor plot analysis script (https://github.com/LAG-MNG-CambridgeUniversity/TCSPCPhasor), from which fluorescence lifetime maps and values were generated. Fluorescence lifetimes are presented as those from individually segmented condensates from 9 images (giving total a total of 86—209 condensates analysed per sample) taken over 3 fully independent experiments. Statistical analysis was performed on Prism 6 (GraphPad, San Diego, CA, USA), where a one-way ANOVA test with Holm-Sidak’s multiple comparison was applied.

### Infrared Nanospectroscopy (AFM-IR)

A nanoIR3 platform (Bruker) combining high resolution and low-noise AFM with a tunable quantum cascade laser (QCL) with top illumination configuration was used. The sample morphology was scanned by the nanoIR3 system, with a line rate within 0.1-0.4 Hz and in contact mode. A silicon gold coated probe with a nominal radius of 30 nm and a cantilever with an elastic constant of about 0.2 N m^−1^ was used. Both infrared (IR) spectra and maps were acquired by using phase loop (PLL) tracking of contact resonance, the phase was zeroed to the desired off-resonant frequency on the left of the IR amplitude maximum and tracked with an integral gain I=0.1-5 and proportional gain P=1-5. All images were acquired with a resolution above 500×100 pixels.

The AFM images were treated and analysed using SPIP software. The height images were first order flattened, while IR and stiffness related maps were only flattened by a zero-order algorithm (offset). Nanoscale-localised spectra were collected by placing the AFM tip on the top of the condensates with a laser wavelength sampling of 2 cm^−1^ and a spectral speed of 100 cm^−1^/s within the range 1462-1800 cm^−1^. Within a single condensate, the spectra were acquired at multiple nanoscale localised positions, the spectrum at each position being the co-average of 5 spectra.

Successively, the spectra were treated by OriginPRO. They were smoothed by an adjacent averaging filter (5 pts) and a Savitzky-Golay filter (second order, 7 points) and normalised. Spectra second derivatives were calculated, smoothed by a Savitzky-Golay filter (second order, 5 points). Relative secondary and quaternary organisation was evaluated by integrating the area of the different secondary structural contributions in the amide band I.

The spectra from 3 different condensates (FUS, n>100; FUS+DNAJB6, n>80) were averaged and used to determine the secondary structure of the condensates. The error in the determination of the relative secondary structure was calculated over the average of at least 5 independent spectra and it is < ±3%.

Spectra were analysed using the microscope’s built-in Analysis Studio (Bruker) and OriginPRO (OriginLab). All measurements were performed at room temperature, with laser power <2 mW and under controlled Nitrogen atmosphere with residual real humidity below 5%.

### Mammalian Cell Lines and Cell Culture

HEK293T cells used in this study were obtained from ATCC. Cells were maintained at 37°C in a humidified atmosphere with 5% CO_2_. HEK293T cells were grown in Dulbecco’s Modified Eagle Medium (DMEM, Invitrogen) which was supplemented with 10% fetal bovine serum (Gibco) and penicillin-streptomycin (Invitrogen).

### Mammalian Transfection

24 h prior to transfection, 293T cells were seeded at 40-60% confluency into 24-well plates coated for 30 min with a 0.1 mg/mL solution of poly-D-lysine (MP Biomedicals Inc.) and washed with PBS (Gibco) once prior to media and subsequent HEK293T cell addition. The next day, expression plasmid was incubated with Opti-MEM (Gibco) and Lipofectamine 2000 (Invitrogen) for 30 min at room temperature prior to addition to cells, per manufacturer protocol. 20 h after transfection, media was changed. Cells were harvested for protein extraction and western blotting 48 h after transfection.

### Deep Mutational Scanning

Deep mutational scanning libraries were prepared in biological duplicates, wherein PCR mutagenesis, construction of bacterial libraries, and construction of yeast libraries were completed as independent replicates. The DNAJB6 yeast expression vector, pAG413GAL-DNAJB6 was miniprepped immediately prior to use. Variant versions of DNAJB6 were made by a single primer site-directed mutagenesis protocol. Oligos were designed to introduce a degenerate codon, NNK, at each amino acid position. Additionally, each oligo was designed to introduce 2-4 synonymous mutations at the codon immediately prior to the degenerate codon to increase sampling diversity. For each codon, an individual mutagenesis single primer PCR reaction was conducted in technical duplicates. The following PCR mix was used for mutagenesis: 5 μL 5X Q5 reaction buffer, 0.5 μL 10 mM dNTPs, 150 ng DNA, 1.25 μL 10 μM primer, and H2O to 25 μL. The following cycling conditions were used: 1. 98°C, 45 s, 2. 98°C, 15 s, 3. 60°C, 15 s, 4. 72°C, 260 s, 5. Return to step 2 29X, 6. 72°C, 240 s. After PCR, the unmodified backbone was digested with 1 μL DpnI at 37°C for 1 h. After digestion, independent PCR replicates were pooled and sets corresponding to 14 contiguous amino acids were pooled together to enable data analysis using short read Illumina sequencing. 8 total sets were created encompassing 112 mutagenized amino acids. The combined sets were column purified with the Zymo DNA Clean & Concentrator Kit. Each biological replicate of each set was transformed into electrocompetent 10-beta *E. Coli* (New England Biolabs) in triplicate according to manufacturer instructions. Cells were plated following outgrowth and recovery on 15 cm LB agar plates containing ampicillin and colonies were allowed to form for 24 h at 30°C. An individual typical transformation yielded 3-10 million colonies for a total of approximately 9-30 million colonies for each biological replicate of each set. As a quality control measure, 20 colonies from each set were sequenced to ensure editing. All colonies were scraped off plates and plasmid libraries were purified by Midiprep (Zymo). Plasmid libraries were transformed into BY4741 containing the expression plasmid pAG416GAL-FUS. Each biological replicate of each set was transformed in 96 separate transformation reactions to ensure appropriate coverage and allowed 48 hours for outgrowth on SC -ura -his plates at 30°C. Yeast libraries were scraped into 10 mL sterilized PBS and frozen in 20% glycerol. For outgrowth, 600 μL of frozen yeast library was inoculated into 6 mL of SC -ura -his for 18 h in triplicate at 30°C with rotation. After inoculation into galactose media, the remaining cells from the overnight cultures were spun down at 4,000 rpm for 5 minutes and the pellets were frozen at −20°C. For each independent culture outgrowth, 12 μL of saturated overnight culture was inoculated into 6 mL of SC -ura -his galactose for 48 h at 30°C with rotation. Each tube was centrifuged at 4,000 rpm for 5 minutes and the supernatant was discarded. Pellets were resuspended in 300 μL of 200 mM LiOAc with 1% SDS and incubated for 15 minutes at 70°C with shaking at 800 rpm. Afterwards, 900 μL 100% ethanol was added, tubes were vortexed, and centrifuged at 13,000 rpm for 10 minutes. Supernatant was discarded and pellets were allowed to air dry for 20 minutes under flame. Pellets were then resuspended in 200 μL TE and incubated at 42°C for 20 minutes and then centrifuged at 13,000 rpm for 10 minutes. The supernatant containing DNA was collected and stored for further use. Each sample was then independently amplified and subsequently indexed for sequencing on a NextSeq 500/550 (Illumina). Depending on the set, amplification of the mutagenized region was done in 8 technical replicates that were pooled after amplification. The following mix was used for all PCR reactions: 5 μL 5X Q5 reaction buffer, 0.5 μL 10 mM dNTPs, 0.5 μL DNA, 0.125 μL 100 μM forward primer, 0.125 μL 100 μM reverse primer, 0.25 μL Q5 polymerase, and 18.5 μL H2O. Amplification was done for 24 cycles for non-induced libraries and 28 cycles for induced samples with the following conditions: 1. 98°C, 45 s, 2. 98°C, 15 s, 3. 58°C, 15 s, 4. 72°C, 30 s, 5. Return to step 2, either 24x or 28x 72°C, 240 s. After pooling, index sequences were attached using the same PCR mix and cycling conditions, but for 8 cycles of amplification. Amplicons were pooled according to set and amplicon length and gel purified to remove primers.

Sequencing data were processed on Illumina Basespace according to default QC settings and downloaded as fastq files. Sequences were aligned using custom Python code. A raw activity score was calculated as:

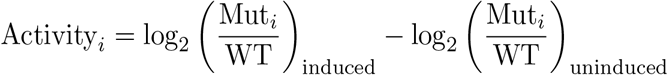

where the subscript i denotes individual unique variants. Muti denotes the average number of counts of the particular mutant codon of interest, averaged over all codings, while WT denotes the number of counts of the wild-type nucleotide sequence. Raw activity scores were then normalized across sets by anchoring stop codon mutants to −1 and WT to 0 to eliminate set by set variation that may have arisen due to experimental fluctuation. The final normalized activity scores are presented in heatmap format for easy visualization. Plotted “wild-type” values are derived from the recoded versions of the wild-type residue at a given position and thus do not always have a 0 value. Raw sequencing reads have been uploaded to the NCBI SRA under BioProject PRJNA769721 (SUB10503160).

### Cas9 Knockout of DNAJB6

A Cas9 expressing HEK293T cell line was generated in a well of a 24-well dish by transfecting 300 ng of pB-CAGGS-Cas9-SV40-BPSV40 vector a long with 100 ng of a plasmid expressing the Piggybac transposase (System Biosciences, LLC) with Lipofectamine 2000 according to manufacturer protocols (Invitrogen). Media was changed 24 h after transfection before selecting with Noursethricin N-Acetyl Transferase (NAT) at 300 μg/mL 48 h after transfection. Cells were expanded and continuously selected with NAT for 2 weeks before being frozen down for further use.

A gRNA lentivirus compatible plasmid encoding two guide RNAs from the Brunello library (GCATATGAAGTGCTGTCGGA and GACTTCTTTGGGAATCGAAG) targeting DNAJB6 was created. Lentivirus was created from this plasmid by co-transfecting HEK293T cells with this plasmid alongside psPAX2 (Addgene #12260) and MD2.G (Addgene #12259). After transfection and a media change 24 h after transfection, media containing lentivirus was harvested 72 h later. Lentivirus containing media was added to Cas9 containing cells for 24 h before a media change. Beginning 48 h after the media change, cells were exposed to 2 weeks of alternating drug selections of NAT at 500 μg/mL and Blasticidin at 2 μg/mL every 48 h with regular splitting, to ensure both Cas9 and gRNA maintained good expression and were not silenced during the outgrowth process. After 2 weeks, single cells were sorted into 96 well plates using the Bigfoot Spectral Cell Sorter (Thermo). To gate on single cells, forward scatter and side scatter were used to isolate single cells and sort them into 100 μL of media. Single cells were allowed to expand for 2 weeks under alternating drug selection. DNA was harvested with QuickExtract (Lucigen) according to manufacturer protocols. PCR primers spanning individual cut sites in addition to primers spanning a potential deletion were used to amplify out the region of DNAJB6 subject to cutting. PCR products were sanger sequenced and TIDE was used to analyze sanger fragments to confirm disruption of DNAJB6^77^. To confirm loss of DNAJB6, western blots were performed. The same process was repeated with non-targeting control guide RNAs (AAAAAGCTTCCGCCTGATGG and AAAACAGGACGATGTGCGGC).

### RNA-seq

HEK293T cells were transfected as previously described with 50 ng of expression plasmid and grown for 72 hours in a 24 well dish after transfection. Cells were harvested in TRIzol and stored at −80°C. RNA was harvested from cells with the Direct-zol miniprep kit (Zymo). Harvested RNA was prepared for sequencing with the NEBNext® Ultra™ II RNA Library Prep Kit for Illumina (NEB). Two biological replicates were performed for each condition. Each individual replicate was amplified with a unique combination of indexing primers after the adaptor ligation step to uniquely identify it. Pooled libraries were combined and sequenced with a 75 cycles NextSeq 500/550 High Output Kit on a NextSeq 550 machine (Illumina). Each replicate was allocated ~30 million reads. Reads were aligned to the hg19 genome using HISAT2 to obtain counts. Differential expression was calculated using limma. Raw sequencing reads have been uploaded to the NCBI SRA under BioProject PRJNA769721 (SUB10426285).

### Human ORFeome Library construction

The pooled hORFeome V8.1 library was inserted into the pAG413GAL-ccdb-6Stop vector with Gateway LR II Clonase Enzyme mix (Invitrogen) at a ratio of 150ng:50ng. The reaction was incubated overnight at 25°C. Expression plasmids were electroporated into electrocompetent 10-beta *E. Coli* (NEB) and ~200,000 colonies were harvested and Miniprepped. The expression plasmid library was transformed into BY4742 and selected in SC -his glucose plates for 48 hours. Individual colonies were picked and arrayed into 96 well plates and saved. To identify the ORF present in each well, each plate was process individually. A total of 20 pools per plate were made, consisting of 12 column pools and 8 row pools. DNA from each of these pools was then obtained using a LiOAc-based extraction. ORFs within each pool were amplified for 30 cycles with general primers binding to the galactose promoter and cyc terminator and subsequently column purified. 250 ng of the purified PCR product was processed with the NEBNext Ultra II FS DNA Library Prep Kit for Illumina (NEB) according to manufacturer protocols. After adaptor ligation and prior to indexing, a forward primer placed 60 bp upstream from the ATG start codon was used in combination with an adaptor reverse primer and amplified for 13 cycles to selectively enrich for the human ORF containing fragment in preparation for sequencing. Each individual pool was then uniquely indexed and sequenced with a 150 cycles NextSeq 500/550 High Output Kit on a NextSeq 500/550 machine (Illumina). After sequencing, the identity of each well was determined by its presence in a specific combination of row and column wells.

## Acknowledgements

We thank members of the Chavez lab for helpful discussions and insights regarding the project. Aaron Gitler provided initial insight and a number of yeast disease models, for which we are immensely grateful. We thank the ALS Stem Cell Core Program at Columbia University Irving Medical Center and its members, Hynek Wichterle, Jon Costa, Emily Lowry, and Niraj Ramsamooj for helpful discussions regarding cell-based assays and for valuable reagents. Vlad Korobeynikov provided helpful guidance and insight into western blotting and protein extraction from cells. Matt Harms provided helpful discussion regarding DNAJB6 patient mutations and insight into interpretation of DMS results. Edwin Chan provided the SoxN plasmids. Richard Gardner provided the PAB1 poly-alanine plasmids. Priya Banerjee provided insight into the phase separation properties of DNAJB6 and FUS. Divyansh Argawal provided helpful discussions regarding statistical analysis of the screening platform. Debbie Hong and Sho Iketani provided valuable guidance on how to perform our deep mutational scanning studies. A.C. is supported by a Career Awards for Medical Scientists from the Burroughs Wellcome Fund and a Therapeutic Idea Award (#AL190073) from the DoD, and a fellowship award to the laboratory from Project ALS. S.J.R is supported by NIH grant F31NS111851. J.W. is supported by NSF grant 2113646. N.Z. is supported by NIH grant 2R01HG006137-10.

## Author contributions

A.C. conceived the project. A.C., S.J.R., S.Q., N.S., and P.S.G. planned and designed experiments. S.J.R. and A.C. performed yeast multiplexed screening. S.J.R. and L.H.H. performed secondary validation testing. S.J.R. and J.S. performed and conducted analysis on deep mutational scanning approaches. S.J.R. and S.M. made cell lines. S.J.R. performed mammalian cell culture assays. S.Q. and J.N. performed *in vitro* studies with DNAJB6 and FUS. C.W.C, C.F.K, and G.S.K.S. performed FLIM experiments. X.C and F.S.R performed AFM-IR experiments. S.J.R., N.Z., and J.W. designed and performed analysis. S.J.R. and A.C. wrote the manuscript with input from all authors.

## Data and materials availability

All reagents generated in this study will be deposited to Addgene. Code used for analysis of the screening approach is available at the Chavez group github account (https://github.com/ChavezResearchLab).

## References

1 Hebert, L. E., Weuve, J., Scherr, P. A. & Evans, D. A. Alzheimer disease in the United States (2010-2050) estimated using the 2010 census. Neurology 80, 1778–1783, doi:10.1212/WNL.0b013e31828726f5 (2013).

2 Vanni, S., Colini Baldeschi, A., Zattoni, M. & Legname, G. Brain aging: A Ianus-faced player between health and neurodegeneration. J Neurosci Res 98, 299–311, doi:10.1002/jnr.24379 (2020).

3 Cavazzoni, P. The Path Forward: Advancing Treatments and Cures for Neurodegenerative Diseases, <https://www.fda.gov/news-events/congressional-testimony/path-forward-advancing-treatments-and-cures-neurodegenerative-diseases-07292021> (2021).

4 Labbadia, J. & Morimoto, R. I. The biology of proteostasis in aging and disease. Annu Rev Biochem 84, 435–464, doi:10.1146/annurev-biochem-060614-033955 (2015).

5 Hipp, M. S., Kasturi, P. & Hartl, F. U. The proteostasis network and its decline in ageing. Nat Rev Mol Cell Biol 20, 421–435, doi:10.1038/s41580-019-0101-y (2019).

6 Outeiro, T. F. & Lindquist, S. Yeast cells provide insight into alpha-synuclein biology and pathobiology. Science 302, 1772–1775, doi:10.1126/science.1090439 (2003).

7 Armakola, M. et al. Inhibition of RNA lariat debranching enzyme suppresses TDP-43 toxicity in ALS disease models. Nat Genet 44, 1302–1309, doi:10.1038/ng.2434 (2012).

8 Sun, Z. et al. Molecular determinants and genetic modifiers of aggregation and toxicity for the ALS disease protein FUS/TLS. PLoS Biol 9, e1000614, doi:10.1371/journal.pbio.1000614 (2011).

9 Gitler, A. D. Beer and bread to brains and beyond: can yeast cells teach us about neurodegenerative disease? Neurosignals 16, 52–62, doi:10.1159/000109759 (2008).

10 Kim, H. J. et al. Therapeutic modulation of eIF2α phosphorylation rescues TDP-43 toxicity in amyotrophic lateral sclerosis disease models. Nat Genet 46, 152–160, doi:10.1038/ng.2853 (2014).

11 Cooper, A. A. et al. Alpha-synuclein blocks ER-Golgi traffic and Rab1 rescues neuron loss in Parkinson’s models. Science 313, 324–328, doi:10.1126/science.1129462 (2006).

12 Couthouis, J. et al. A yeast functional screen predicts new candidate ALS disease genes. Proc Natl Acad Sci U S A 108, 20881–20890, doi:10.1073/pnas.1109434108 (2011).

13 Kryndushkin, D., Ihrke, G., Piermartiri, T. C. & Shewmaker, F. A yeast model of optineurin proteinopathy reveals a unique aggregation pattern associated with cellular toxicity. Mol Microbiol 86, 1531–1547, doi:10.1111/mmi.12075 (2012).

14 Treusch, S. et al. Functional links between Aβ toxicity, endocytic trafficking, and Alzheimer’s disease risk factors in yeast. Science 334, 1241–1245, doi:10.1126/science.1213210 (2011).

15 Ju, S. et al. A yeast model of FUS/TLS-dependent cytotoxicity. PLoS Biol 9, e1001052, doi:10.1371/journal.pbio.1001052 (2011).

16 Chen, Y. C. et al. Randomized CRISPR-Cas Transcriptional Perturbation Screening Reveals Protective Genes against Alpha-Synuclein Toxicity. Mol Cell 68, 247–257.e245, doi:10.1016/j.molcel.2017.09.014 (2017).

17 Lee, J. C. et al. Inhibition of p38 MAP kinase as a therapeutic strategy. Immunopharmacology 47, 185–201, doi:10.1016/s0162-3109(00)00206-x (2000).

18 Jo, M. et al. Yeast genetic interaction screen of human genes associated with amyotrophic lateral sclerosis: identification of MAP2K5 kinase as a potential drug target. Genome Res 27, 1487–1500, doi:10.1101/gr.211649.116 (2017).

19 Guerrero, E. N. et al. TDP-43/FUS in motor neuron disease: Complexity and challenges. Prog Neurobiol 145-146, 78–97, doi:10.1016/j.pneurobio.2016.09.004 (2016).

20 Yan, Z. et al. Yeast Barcoders: a chemogenomic application of a universal donor-strain collection carrying bar-code identifiers. Nat Methods 5, 719–725, doi:10.1038/nmeth.1231 (2008).

21 Schmierer, B. et al. CRISPR/Cas9 screening using unique molecular identifiers. Mol Syst Biol 13, 945, doi:10.15252/msb.20177834 (2017).

22 Kim, H. J. et al. Mutations in prion-like domains in hnRNPA2B1 and hnRNPA1 cause multisystem proteinopathy and ALS. Nature 495, 467–473, doi:10.1038/nature11922 (2013).

23 Suzuki, H. & Matsuoka, M. The Lysosomal Trafficking Transmembrane Protein 106B Is Linked to Cell Death. J Biol Chem 291, 21448–21460, doi:10.1074/jbc.M116.737171 (2016).

24 Jovičić, A. et al. Modifiers of C9orf72 dipeptide repeat toxicity connect nucleocytoplasmic transport defects to FTD/ALS. Nat Neurosci 18, 1226–1229, doi:10.1038/nn.4085 (2015).

25 Zheng, C., Geetha, T. & Babu, J. R. Failure of ubiquitin proteasome system: risk for neurodegenerative diseases. Neurodegener Dis 14, 161–175, doi:10.1159/000367694 (2014).

26 Arndt, V., Rogon, C. & Höhfeld, J. To be, or not to be--molecular chaperones in protein degradation. Cell Mol Life Sci 64, 2525–2541, doi:10.1007/s00018-007-7188-6 (2007).

27 Jackrel, M. E. et al. Potentiated Hsp104 variants antagonize diverse proteotoxic misfolding events. Cell 156, 170–182, doi:10.1016/j.cell.2013.11.047 (2014).

28 Ioakeimidis, F. et al. A splicing mutation in the novel mitochondrial protein DNAJC11 causes motor neuron pathology associated with cristae disorganization, and lymphoid abnormalities in mice. PLoS One 9, e104237, doi:10.1371/journal.pone.0104237 (2014).

29 Chen, X. & Petranovic, D. Amyloid-β peptide-induced cytotoxicity and mitochondrial dysfunction in yeast. FEMS Yeast Res 15, doi:10.1093/femsyr/fov061 (2015).

30 Keefer, K. M., Stein, K. C. & True, H. L. Heterologous prion-forming proteins interact to cross-seed aggregation in Saccharomyces cerevisiae. Scientific Reports 7, 5853, doi:10.1038/s41598-017-05829-5 (2017).

31 Higurashi, T., Hines, J. K., Sahi, C., Aron, R. & Craig, E. A. Specificity of the J-protein Sis1 in the propagation of 3 yeast prions. Proceedings of the National Academy of Sciences 105, 16596–16601, doi:10.1073/pnas.0808934105 (2008).

32 Breslow, D. K. et al. A comprehensive strategy enabling high-resolution functional analysis of the yeast genome. Nat Methods 5, 711–718, doi:10.1038/nmeth.1234 (2008).

33 Park, S. K. et al. Development and validation of a yeast high-throughput screen for inhibitors of Aβ_42_ oligomerization. Dis Model Mech 4, 822–831, doi:10.1242/dmm.007963 (2011).

34 Sarnoski, E. A., Liu, P. & Acar, M. A High-Throughput Screen for Yeast Replicative Lifespan Identifies Lifespan-Extending Compounds. Cell Rep 21, 2639–2646, doi:10.1016/j.celrep.2017.11.002 (2017).

35 Wong, L. H. et al. A yeast chemical genetic screen identifies inhibitors of human telomerase. Chem Biol 20, 333–340, doi:10.1016/j.chembiol.2012.12.008 (2013).

36 Zhang, L. et al. A high-throughput screen for chemical inhibitors of exocytic transport in yeast. Chembiochem 11, 1291–1301, doi:10.1002/cbic.200900681 (2010).

37 Kachroo, A. H. et al. Evolution. Systematic humanization of yeast genes reveals conserved functions and genetic modularity. Science 348, 921–925, doi:10.1126/science.aaa0769 (2015).

38 Uhlén, M. et al. Proteomics. Tissue-based map of the human proteome. Science 347, 1260419, doi:10.1126/science.1260419 (2015).

39 Thiruvalluvan, A. et al. DNAJB6, a Key Factor in Neuronal Sensitivity to Amyloidogenesis. Mol Cell 78, 346–358.e349, doi:10.1016/j.molcel.2020.02.022 (2020).

40 Brehme, M. et al. A chaperome subnetwork safeguards proteostasis in aging and neurodegenerative disease. Cell Rep 9, 1135–1150, doi:10.1016/j.celrep.2014.09.042 (2014).

41 Fan, C. Y., Lee, S. & Cyr, D. M. Mechanisms for regulation of Hsp70 function by Hsp40. Cell Stress Chaperones 8, 309–316, doi:10.1379/1466-1268(2003)008<0309:mfrohf>2.0.co;2 (2003).

42 Kampinga, H. H. & Craig, E. A. The HSP70 chaperone machinery: J proteins as drivers of functional specificity. Nat Rev Mol Cell Biol 11, 579–592, doi:10.1038/nrm2941 (2010).

43 Gillis, J. et al. The DNAJB6 and DNAJB8 protein chaperones prevent intracellular aggregation of polyglutamine peptides. J Biol Chem 288, 17225–17237, doi:10.1074/jbc.M112.421685 (2013).

44 Kakkar, V. et al. The S/T-Rich Motif in the DNAJB6 Chaperone Delays Polyglutamine Aggregation and the Onset of Disease in a Mouse Model. Mol Cell 62, 272–283, doi:10.1016/j.molcel.2016.03.017 (2016).

45 Månsson, C. et al. DNAJB6 is a peptide-binding chaperone which can suppress amyloid fibrillation of polyglutamine peptides at substoichiometric molar ratios. Cell Stress Chaperones 19, 227–239, doi:10.1007/s12192-013-0448-5 (2014).

46 Hageman, J. et al. A DNAJB chaperone subfamily with HDAC-dependent activities suppresses toxic protein aggregation. Mol Cell 37, 355–369, doi:10.1016/j.molcel.2010.01.001 (2010).

47 Harms, M. B. et al. Exome sequencing reveals DNAJB6 mutations in dominantly-inherited myopathy. Ann Neurol 71, 407–416, doi:10.1002/ana.22683 (2012).

48 Bengoechea, R., Pittman, S. K., Tuck, E. P., True, H. L. & Weihl, C. C. Myofibrillar disruption and RNA-binding protein aggregation in a mouse model of limb-girdle muscular dystrophy 1D. Hum Mol Genet 24, 6588–6602, doi:10.1093/hmg/ddv363 (2015).

49 Chen, H. J. et al. The heat shock response plays an important role in TDP-43 clearance: evidence for dysfunction in amyotrophic lateral sclerosis. Brain 139, 1417–1432, doi:10.1093/brain/aww028 (2016).

50 Udan-Johns, M. et al. Prion-like nuclear aggregation of TDP-43 during heat shock is regulated by HSP40/70 chaperones. Hum Mol Genet 23, 157–170, doi:10.1093/hmg/ddt408 (2014).

51 Murakami, T. et al. ALS/FTD Mutation-Induced Phase Transition of FUS Liquid Droplets and Reversible Hydrogels into Irreversible Hydrogels Impairs RNP Granule Function. Neuron 88, 678–690, doi:10.1016/j.neuron.2015.10.030 (2015).

52 Khong, A. et al. The Stress Granule Transcriptome Reveals Principles of mRNA Accumulation in Stress Granules. Mol Cell 68, 808–820.e805, doi:10.1016/j.molcel.2017.10.015 (2017).

53 Markmiller, S. et al. Context-Dependent and Disease-Specific Diversity in Protein Interactions within Stress Granules. Cell 172, 590–604 e513, doi:10.1016/j.cell.2017.12.032 (2018).

54 Patel, A. et al. A Liquid-to-Solid Phase Transition of the ALS Protein FUS Accelerated by Disease Mutation. Cell 162, 1066–1077, doi:10.1016/j.cell.2015.07.047 (2015).

55 Qamar, S. et al. FUS Phase Separation Is Modulated by a Molecular Chaperone and Methylation of Arginine Cation-π Interactions. Cell 173, 720–734.e715, doi:10.1016/j.cell.2018.03.056 (2018).

56 Krainer, G. et al. Reentrant liquid condensate phase of proteins is stabilized by hydrophobic and non-ionic interactions. Nat Commun 12, 1085, doi:10.1038/s41467-021-21181-9 (2021).

57 Itzhak, D. N., Tyanova, S., Cox, J. & Borner, G. H. Global, quantitative and dynamic mapping of protein subcellular localization. Elife 5, doi:10.7554/eLife.16950 (2016).

58 Kato, M. et al. Cell-free formation of RNA granules: low complexity sequence domains form dynamic fibers within hydrogels. Cell 149, 753–767, doi:10.1016/j.cell.2012.04.017 (2012).

59 Ruggeri, F. S. et al. Infrared nanospectroscopy characterization of oligomeric and fibrillar aggregates during amyloid formation. Nat Commun 6, 7831, doi:10.1038/ncomms8831 (2015).

60 Ruggeri, F. S. et al. Influence of the β-sheet content on the mechanical properties of aggregates during amyloid fibrillization. Angew Chem Int Ed Engl 54, 2462–2466, doi:10.1002/anie.201409050 (2015).

61 Shimanovich, U. et al. Silk micrococoons for protein stabilisation and molecular encapsulation. Nat Commun 8, 15902, doi:10.1038/ncomms15902 (2017).

62 Ruggeri, F. S., Mannini, B., Schmid, R., Vendruscolo, M. & Knowles, T. P. J. Single molecule secondary structure determination of proteins through infrared absorption nanospectroscopy. Nat Commun 11, 2945, doi:10.1038/s41467-020-16728-1 (2020).

63 Ruggeri, F. S. et al. The Influence of Pathogenic Mutations in α-Synuclein on Biophysical and Structural Characteristics of Amyloid Fibrils. ACS Nano 14, 5213–5222, doi:10.1021/acsnano.9b09676 (2020).

64 Harmon, T. S., Holehouse, A. S., Rosen, M. K. & Pappu, R. V. Intrinsically disordered linkers determine the interplay between phase separation and gelation in multivalent proteins. Elife 6, doi:10.7554/eLife.30294 (2017).

65 Gu, J. et al. Hsp40 proteins phase separate to chaperone the assembly and maintenance of membraneless organelles. Proc Natl Acad Sci U S A 117, 31123–31133, doi:10.1073/pnas.2002437117 (2020).

66 Weston, S. et al. A Yeast Suppressor Screen Used To Identify Mammalian SIRT1 as a Proviral Factor for Middle East Respiratory Syndrome Coronavirus Replication. J Virol 93, doi:10.1128/jvi.00197-19 (2019).

67 Fernández-Acero, T. et al. A yeast-based in vivo bioassay to screen for class I phosphatidylinositol 3-kinase specific inhibitors. J Biomol Screen 17, 1018–1029, doi:10.1177/1087057112450051 (2012).

68 Conicella, A. E., Zerze, G. H., Mittal, J. & Fawzi, N. L. ALS Mutations Disrupt Phase Separation Mediated by α-Helical Structure in the TDP-43 Low-Complexity C-Terminal Domain. Structure 24, 1537–1549, doi:10.1016/j.str.2016.07.007 (2016).

69 Tosatto, L. et al. Single-molecule FRET studies on alpha-synuclein oligomerization of Parkinson’s disease genetically related mutants. Sci Rep 5, 16696, doi:10.1038/srep16696 (2015).

70 López-Erauskin, J. et al. ALS/FTD-Linked Mutation in FUS Suppresses Intra-axonal Protein Synthesis and Drives Disease Without Nuclear Loss-of-Function of FUS. Neuron 100, 816–830.e817, doi:10.1016/j.neuron.2018.09.044 (2018).

71 Thelen, M. P. & Kye, M. J. The Role of RNA Binding Proteins for Local mRNA Translation: Implications in Neurological Disorders. Front Mol Biosci 6, 161, doi:10.3389/fmolb.2019.00161 (2019).

72 Birsa, N. et al. FUS-ALS mutants alter FMRP phase separation equilibrium and impair protein translation. Sci Adv 7, doi:10.1126/sciadv.abf8660 (2021).

73 Yu, H. et al. HSP70 chaperones RNA-free TDP-43 into anisotropic intranuclear liquid spherical shells. Science 371, doi:10.1126/science.abb4309 (2021).

74 Aprile, F. A. et al. The molecular chaperones DNAJB6 and Hsp70 cooperate to suppress α-synuclein aggregation. Sci Rep 7, 9039, doi:10.1038/s41598-017-08324-z (2017).

75 Månsson, C. et al. Interaction of the molecular chaperone DNAJB6 with growing amyloid-beta 42 (Aβ42) aggregates leads to sub-stoichiometric inhibition of amyloid formation. J Biol Chem 289, 31066–31076, doi:10.1074/jbc.M114.595124 (2014).

76 Li, W. et al. MAGeCK enables robust identification of essential genes from genome-scale CRISPR/Cas9 knockout screens. Genome Biol 15, 554, doi:10.1186/s13059-014-0554-4 (2014).

77 Brinkman, E. K., Chen, T., Amendola, M. & van Steensel, B. Easy quantitative assessment of genome editing by sequence trace decomposition. Nucleic Acids Res 42, e168, doi:10.1093/nar/gku936 (2014).

78 Sondheimer, N., Lopez, N., Craig, E. A. & Lindquist, S. The role of Sis1 in the maintenance of the [RNQ+] prion. Embo j 20, 2435–2442, doi:10.1093/emboj/20.10.2435 (2001).

79 Bagriantsev, S. N., Gracheva, E. O., Richmond, J. E. & Liebman, S. W. Variant-specific [PSI+] infection is transmitted by Sup35 polymers within [PSI+] aggregates with heterogeneous protein composition. Mol Biol Cell 19, 2433–2443, doi:10.1091/mbc.e08-01-0078 (2008).

80 Barlow, J. T., Bogatyrev, S. R. & Ismagilov, R. F. A quantitative sequencing framework for absolute abundance measurements of mucosal and lumenal microbial communities. Nat Commun 11, 2590, doi:10.1038/s41467-020-16224-6 (2020).

81 McIntyre, L. M. et al. RNA-seq: technical variability and sampling. BMC Genomics 12, 293, doi:10.1186/1471-2164-12-293 (2011).

