## Supplemental Figures and Notes for "A multiplex platform to identify mechanisms and modulators of proteotoxicity in neurodegeneration"

### Supplementary Figures

**Supplementary Figure 1. Detailed development of a multiplexed screening platform. a.** Individual yeast strains containing an integrated DNA barcode are transformed with a construct encoding an aggregation-prone protein associated with neurodegeneration before pooling. **b.** Rescuers are introduced *en masse* through mating and selection. **c.** Mated barcode pools are grown in inducing media in 96-well plate format before DNA harvesting, NGS, and subsequent analysis.

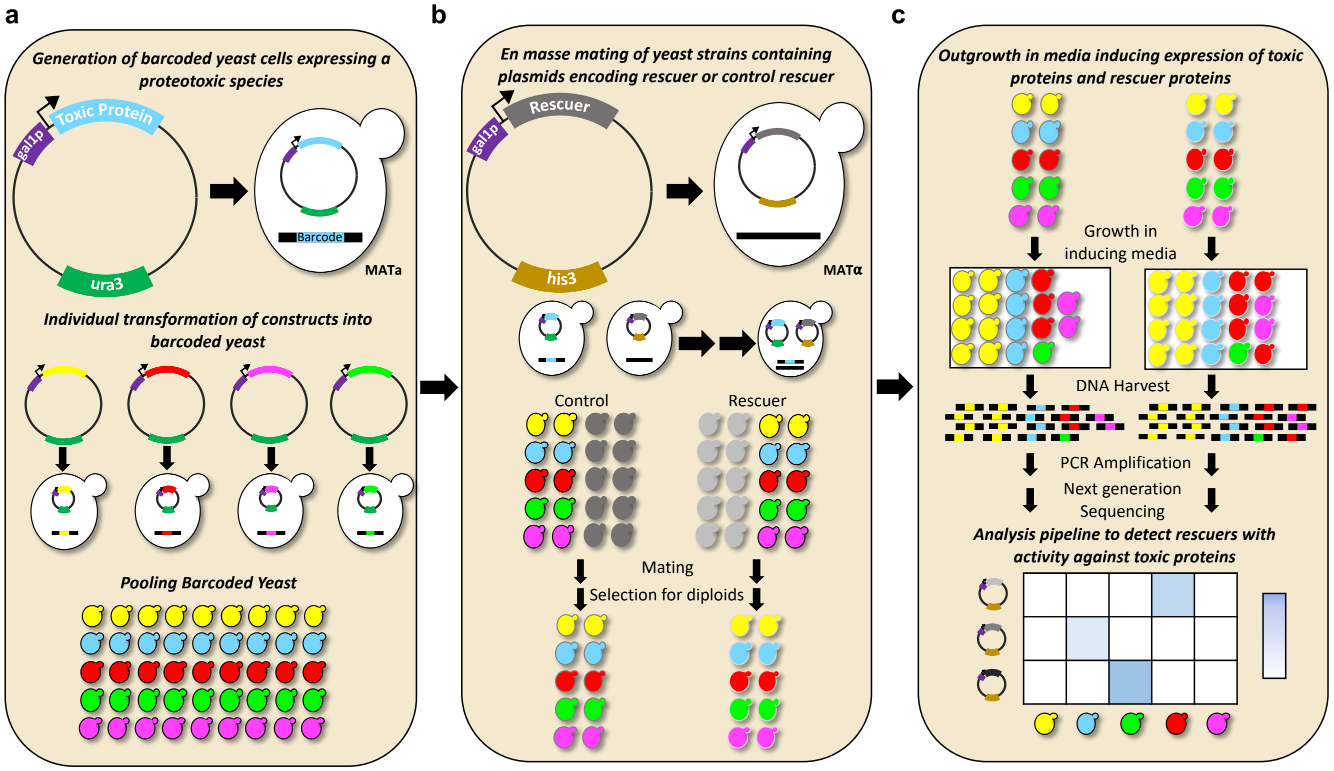

**Supplementary Figure 2.** ***En masse*** **mating of a barcode pool and outgrowth of a mated barcoded pool do not perturb barcode ratios. a.** Example of correlation plot between two separately mated pools that have been selected for diploids, each dot represents a different barcode within the population. **b.** Correlation values for 36 comparisons between the barcode abundance for separately mated pools. **c.** Example of correlation plot between two separately mated pools that have been selected for diploids, and outgrown in inducing media, each dot represents a different barcode within the population. **d.** Correlation values for 36 comparisons between separately mated and outgrown pools.

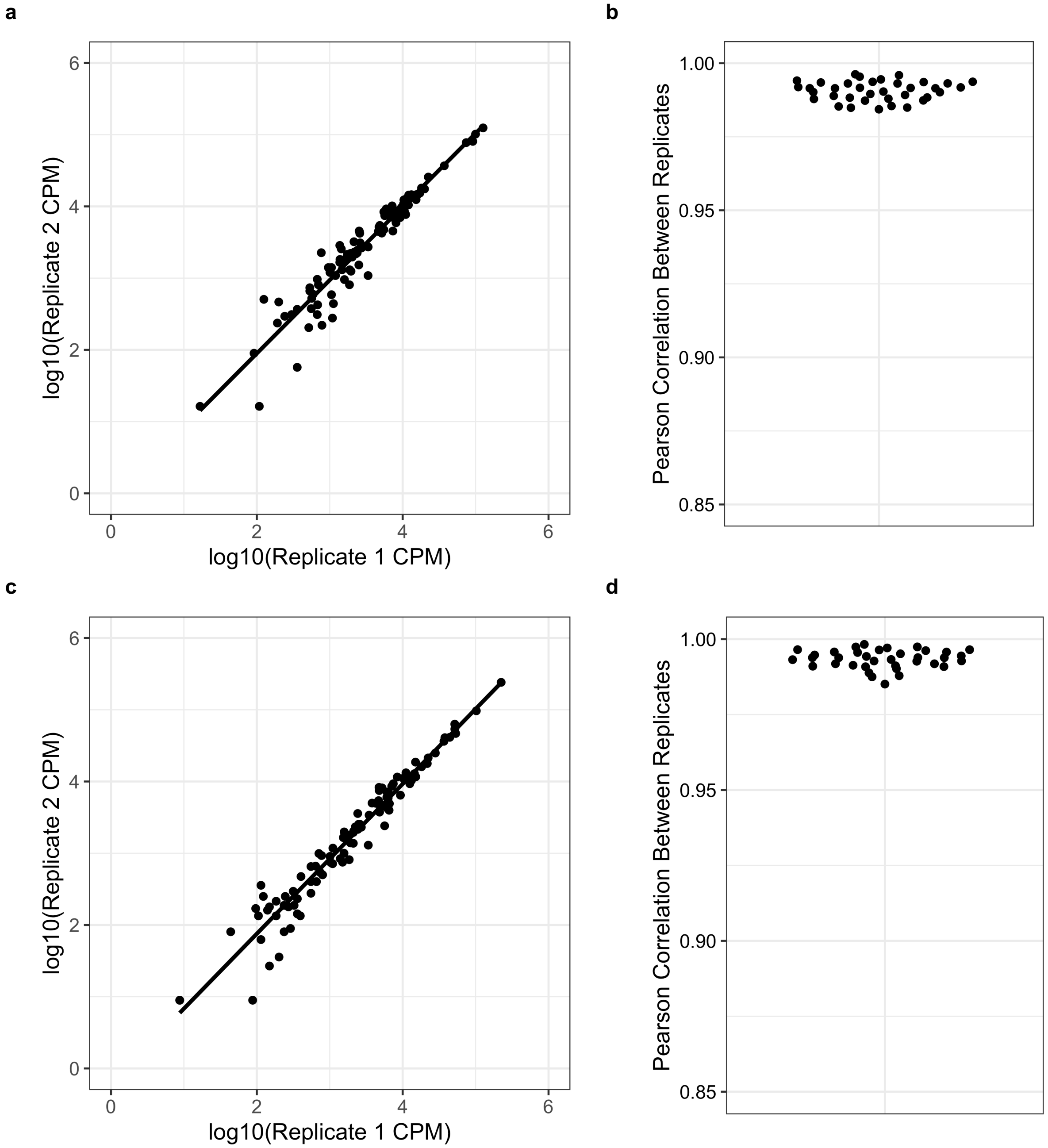

**Supplementary Figure 3. Comparison of three pooling strategies for detecting known interactions**. **a.** Log2 fold change heatmaps for 3 pooling strategies. Previously known interactions that are expected are outlined in purple **b.** Correlations between barcodes after pooled barcoded strains were mated to the same control rescuer, selected for diploids, and grown under inducing condition using each of the three different pooling strategies. **c.** Coefficient of variation vs. relative barcode abundance plot for each of the 3 pooling strategies.

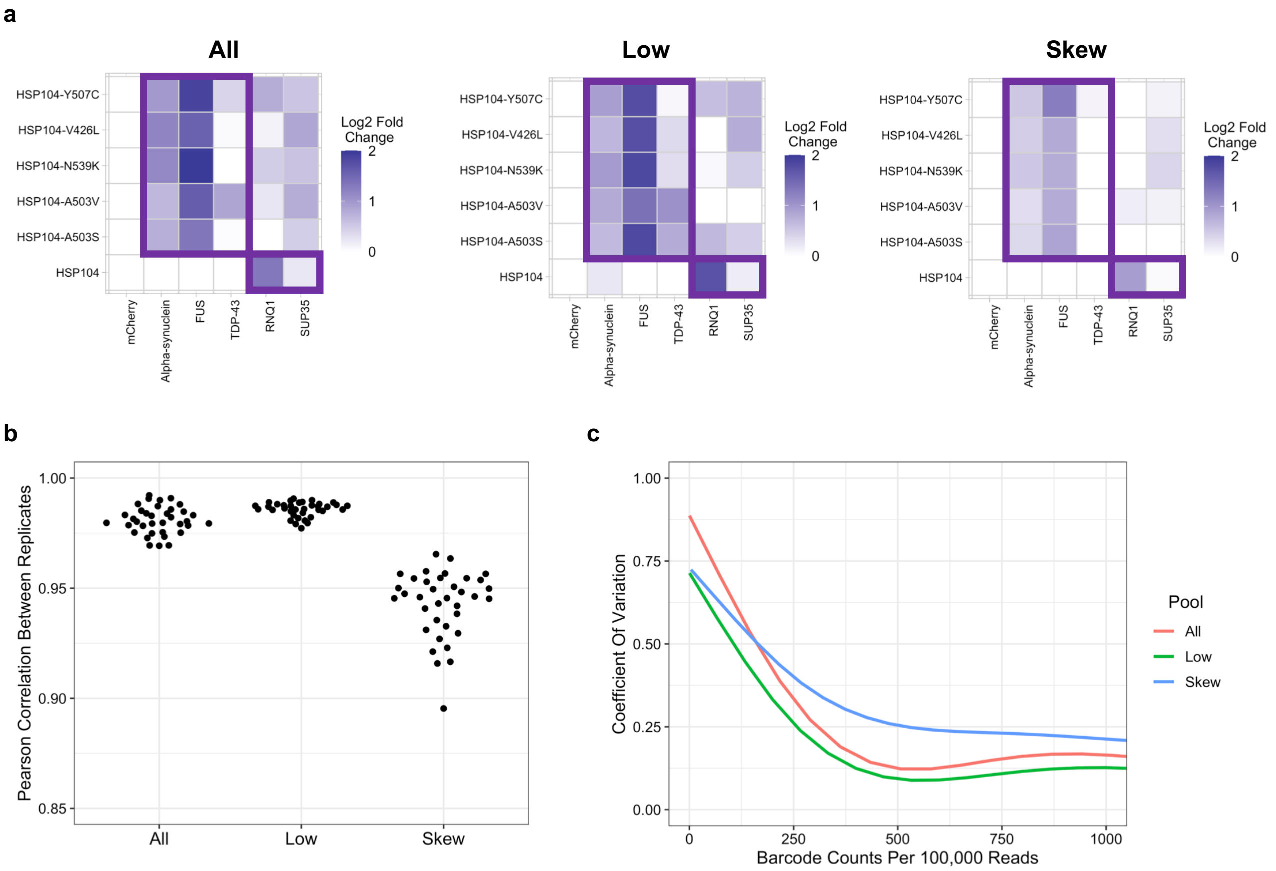

**Supplementary Figure 4. Exploration of biological and technical sources of error for the optimal screening strategy. a.** Correlation between biological replicates (separately mated, selected, outgrown, harvested, and PCR amplified) and technical replicates (same sample of harvested DNA separately PCR amplified). **b.** Coefficient of variation vs. relative barcode abundance plot for biological replicates of pooled DNA-barcoded library mated to an inert rescuer demonstrating the effect of averaging between biological replicates. **c.** Coefficient of variation vs. relative barcode abundance plot for technical replicates of pooled DNA-barcoded library mated to an inert rescuer demonstrating effect of averaging between technical replicates.

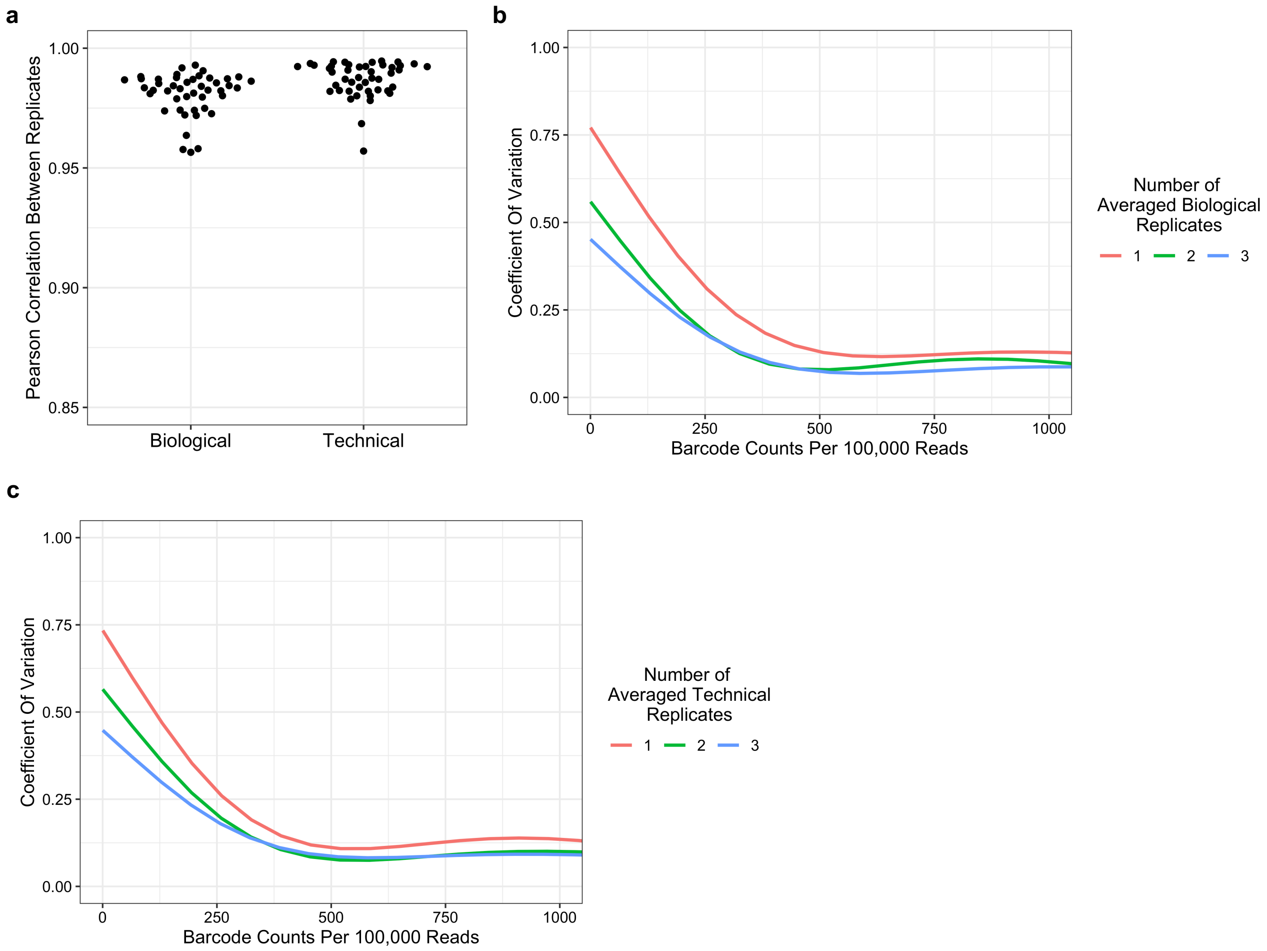

**Supplementary Figure 5. Validation of the optimized multiplexed screening approach using known genetic interactions. a.** Optimized conditions enable detection of positive control interactions in pilot screen **b-g.** Spot assay validation of tested interactions in pilot multiplexed screen.

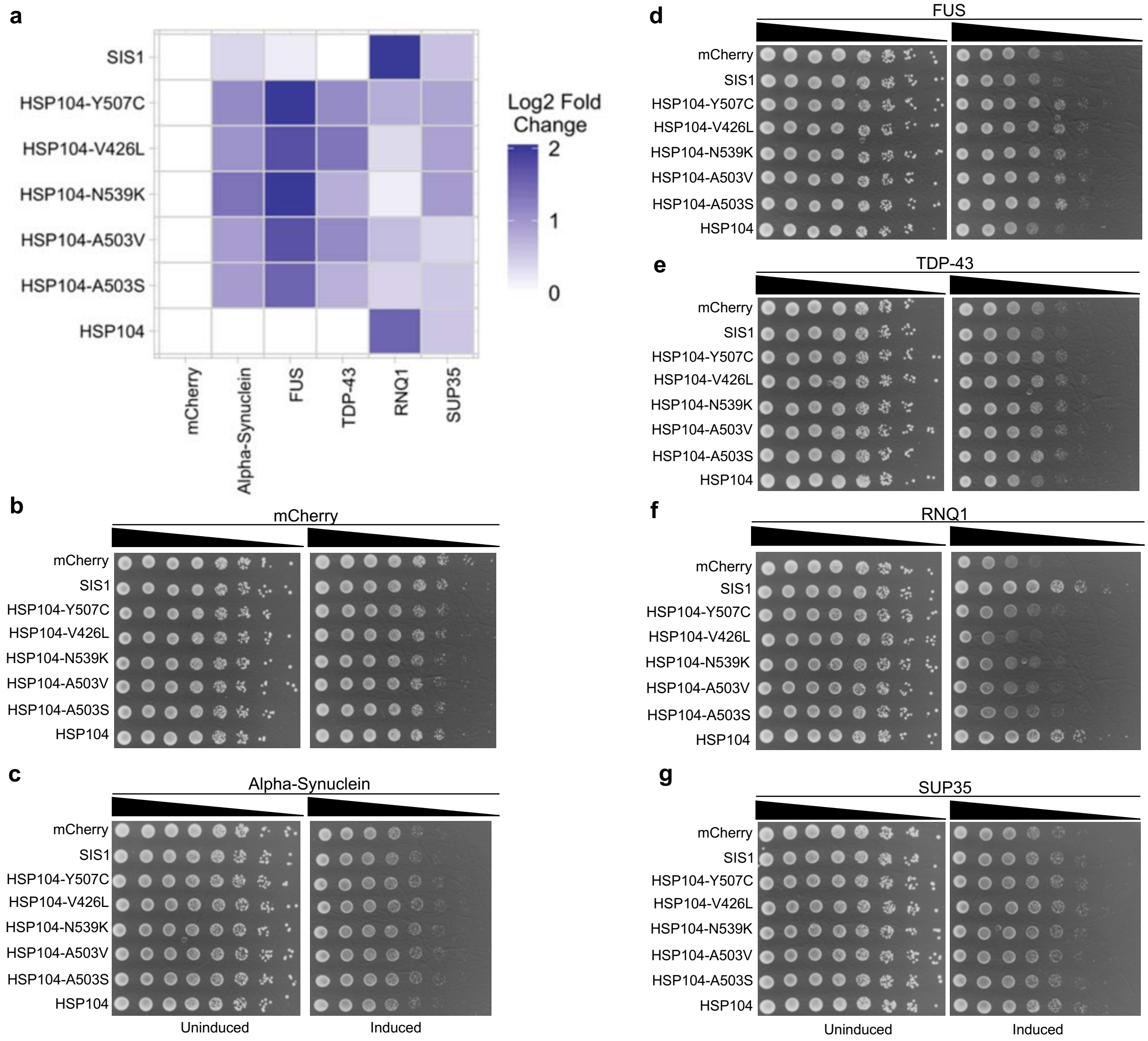

**Supplementary Figure 6. Simulation of how redundant barcoding enhances the ability to reject the null hypothesis.** Fold change required for rejection of null hypothesis simulated using optimized pooling conditions with two biological replicates and two technical replicates.

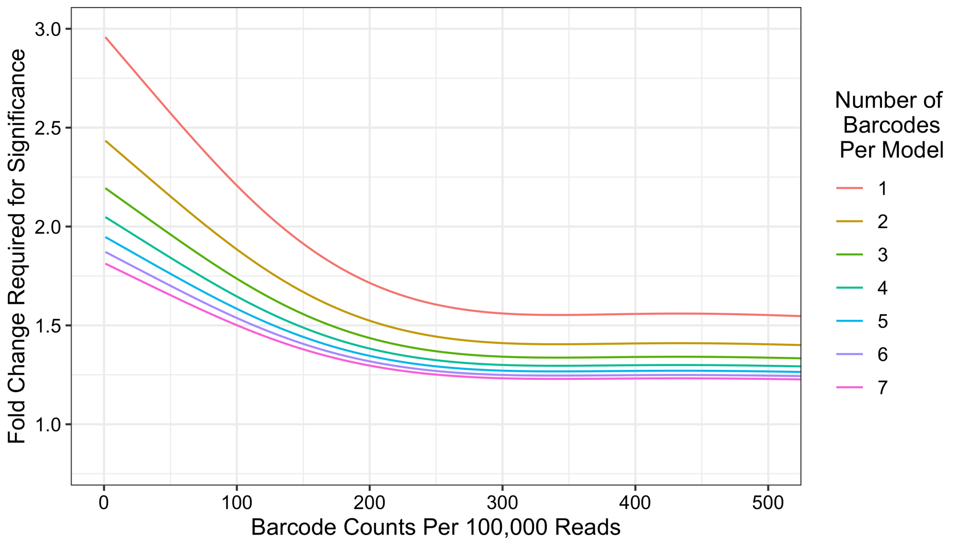

**Supplementary Figure 7. Determination of number of reads required to adequately sample 302-member DNA-barcoded pool**. **a.** Coefficient of variation vs. relative barcode abundance for DNA-barcoded library mated to an inert rescuer at different levels of read subsampling. **b.** Number of individual barcoded strains with less than 10 raw reads at different levels of read subsampling.

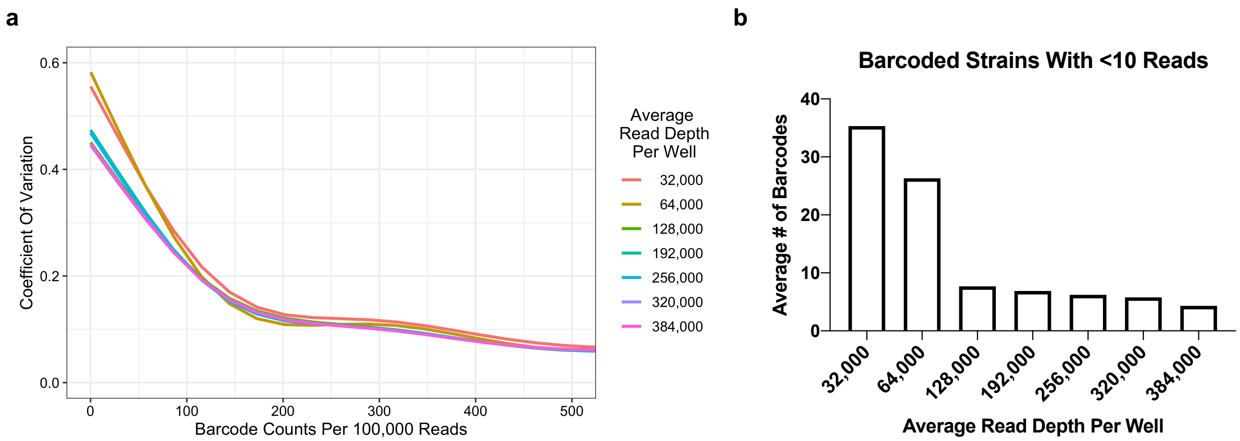

**Supplementary Figure 8. Full yeast chaperone screen. a.** Log2 fold change interactions between all tested yeast chaperones and the models included in the pool. **b.** Significant interactions between yeast chaperones and models included in the pool.

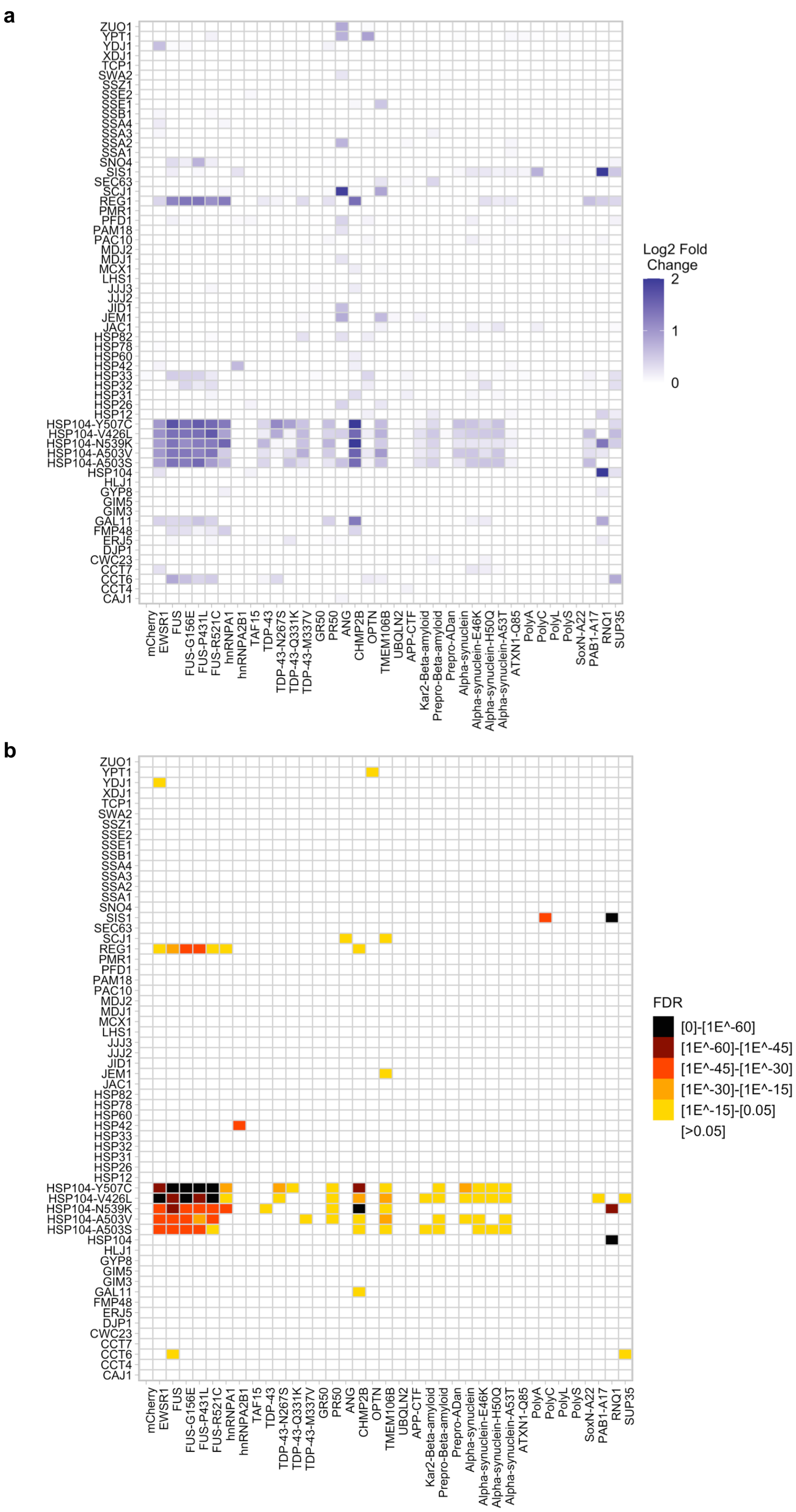

**Supplementary Figure 9. Full human chaperone screen. a.** Log2 fold change interactions between all tested human chaperones and the models included in the pool. **b.** Significant interactions between human chaperones and models included in the pool.

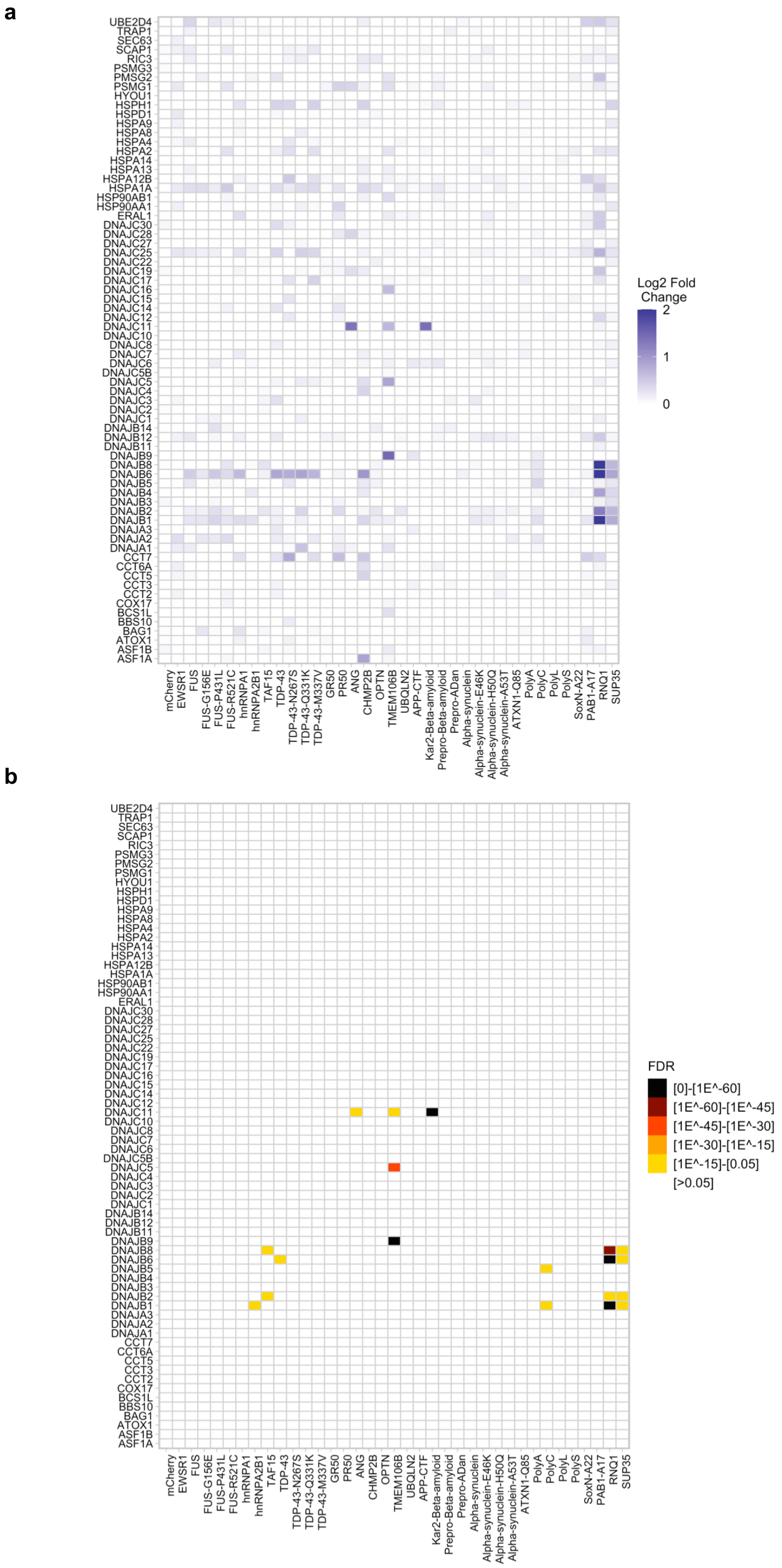

**Supplementary Figure 10. Validation of liquid culture growth assay.** The interactions tested in our pilot screen for **a.** mCherry **b.** Alpha-synuclein **c.** FUS **d.** TDP-43 **e.** RNQ1 and **f.** SUP-35 and validated with spot assays were re-tested with the liquid culture growth assay to validate its behavior. Data are shown as mean ± s.d. for three biological replicates. Comparisons were conducted with ordinary one-way ANOVA with display of comparisons of interactions with in positive changes in relative growth; ns = not significant, *P≤0.05, **P≤0.01, ***P≤0.001, ****P≤0.0001.

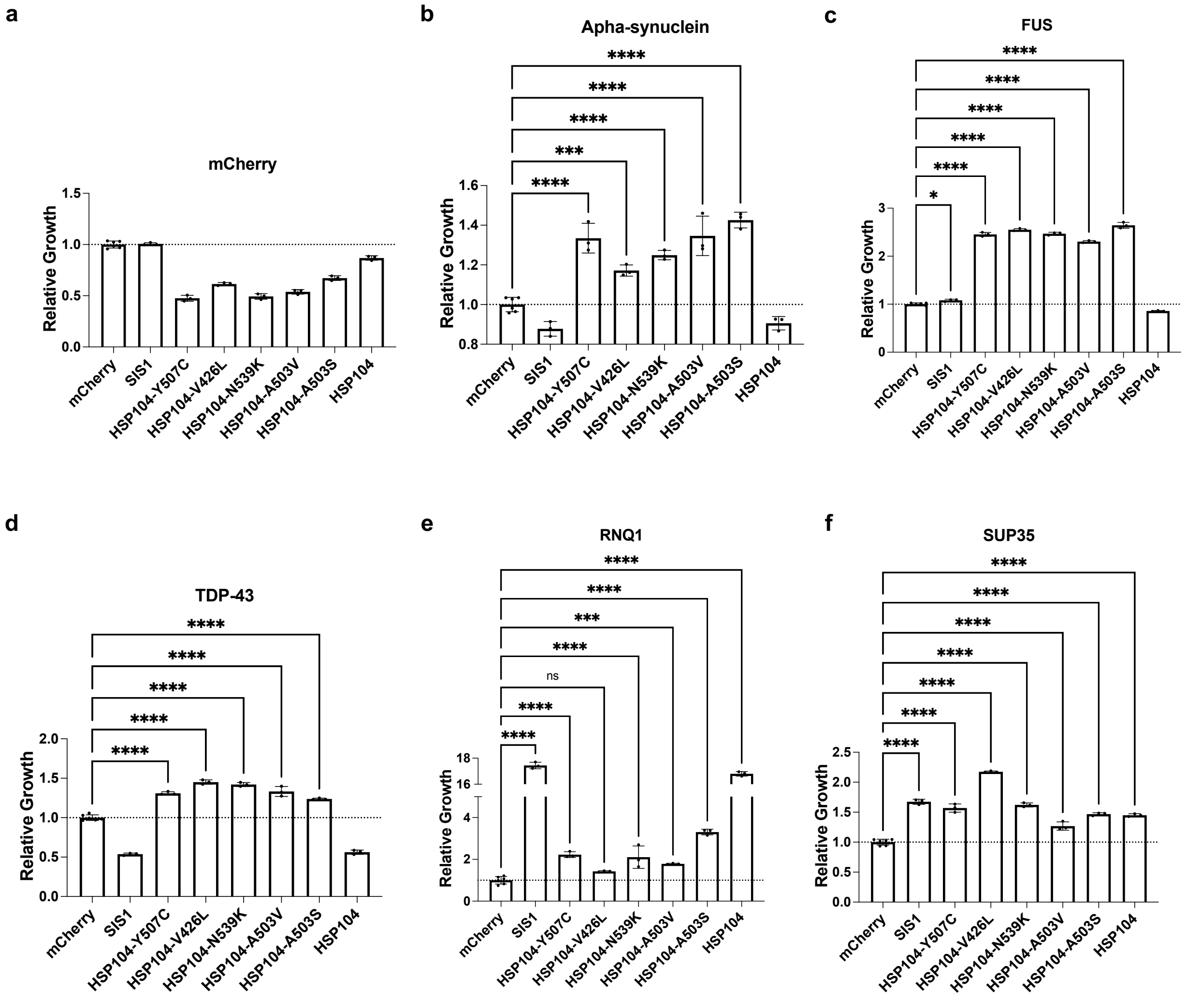

**Supplementary Figure 11. Subsampling of barcodes per model demonstrates power of redundant barcoding.** Yeast and human rescuers with called hits were reanalyzed with fewer number of barcoded strains included in the analysis. All hits shown in the 5-7 Barcodes/Model condition were validated.

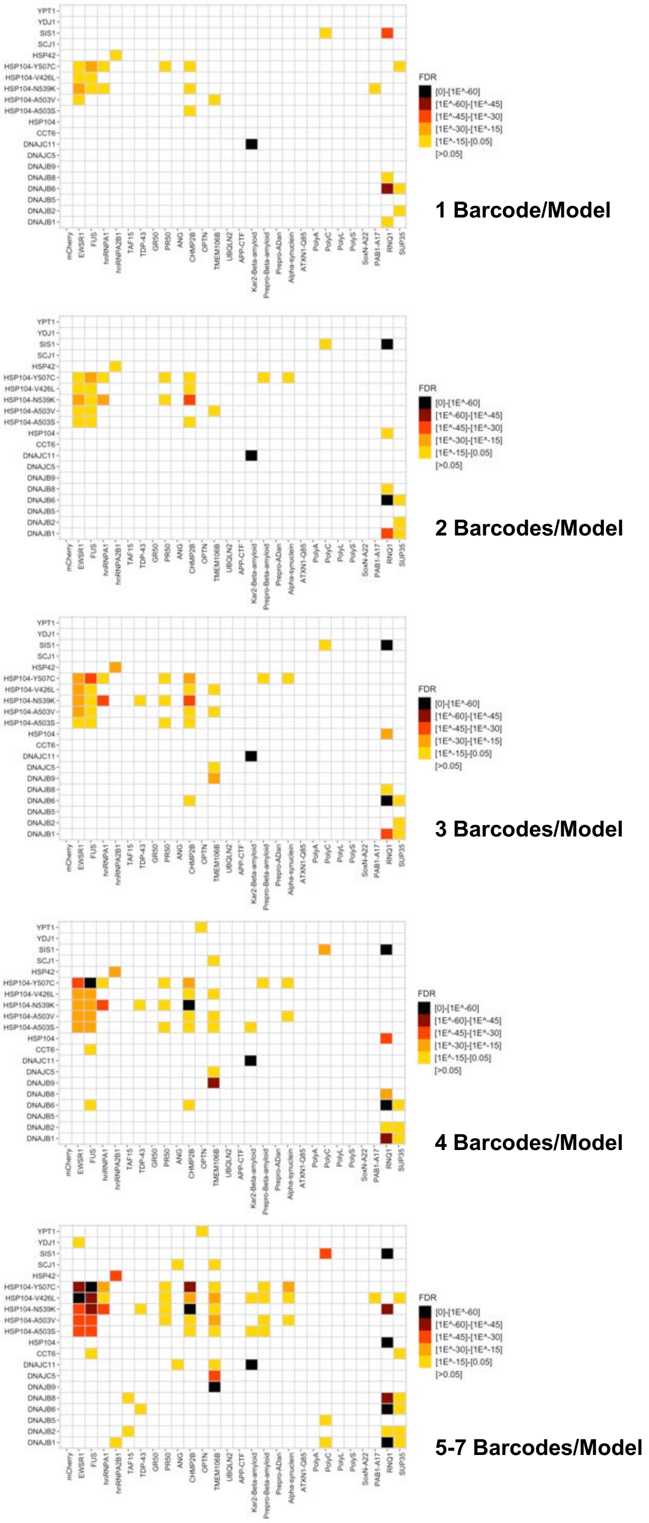

**Supplementary Figure 12. DNAJB6 shows specific activity against FUS, TDP-43, and hnRNPA1. a-c.** DNAJB6 was tested alongside other human HSP40 proteins for their ability to rescue **a.** FUS, **b.** TDP-43, and **c.** hnRNPA1 proteotoxicity in yeast. Comparisons were conducted with ordinary one-way ANOVA with display of comparisons of interactions with in positive changes in relative growth; *P≤0.05, ***P≤0.001, ****P≤0.0001.

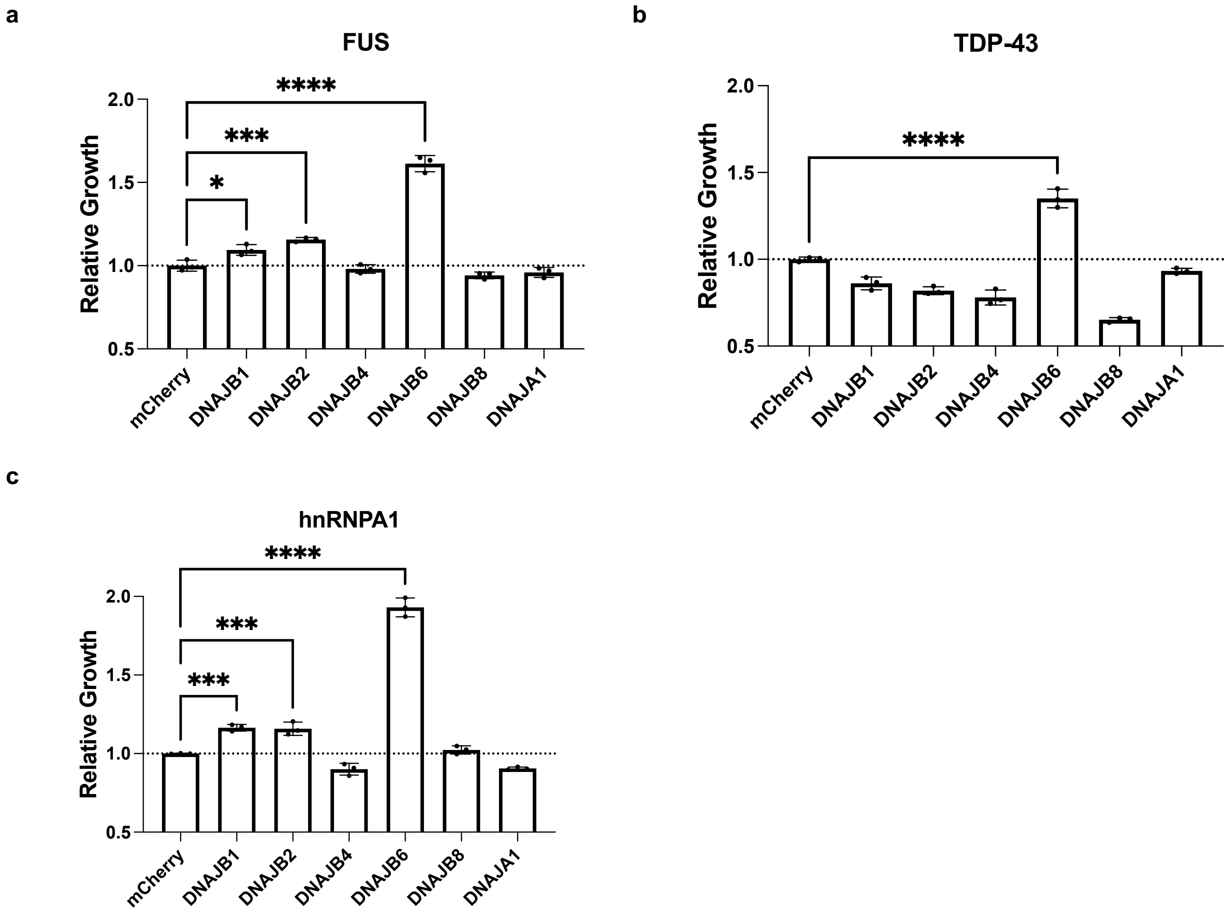

**Supplementary Figure 13. RNA-seq demonstrates that DNAJB6 is significantly upregulated in response to FUS or TDP-43 overexpression in HEK293T cells.** Volcano plots for HEK293T expressed chaperones for **a.** FUS and **b.** TDP-43 compared to EYFP overexpressing cells. Two biological replicates were done for each condition.

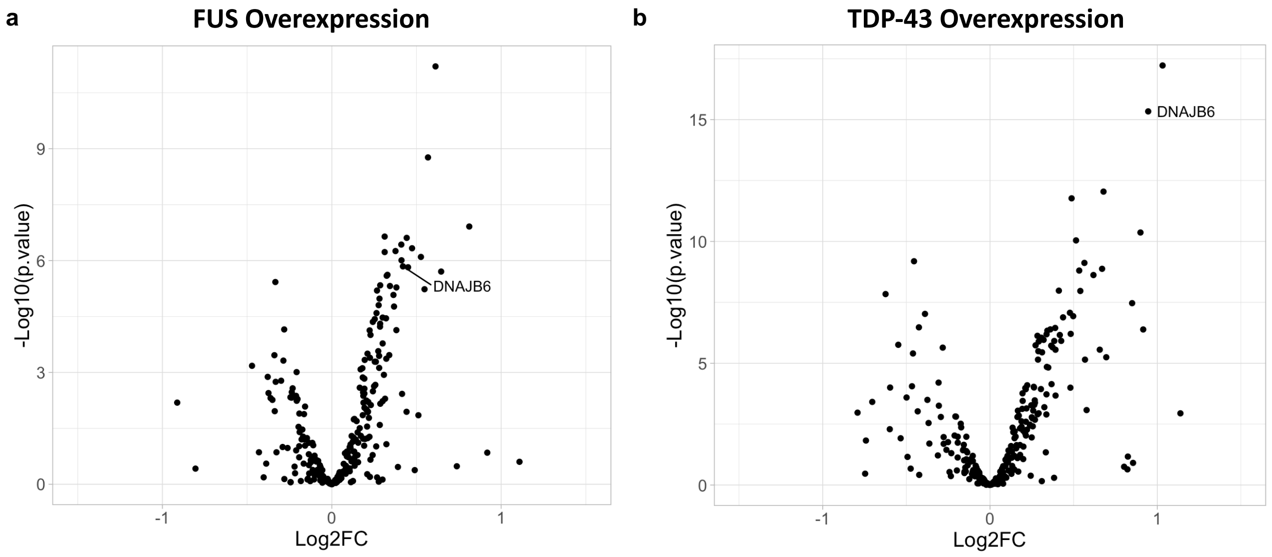

**Supplementary Figure 14. Knockout of DNAJB6 does not impact SDS solubility of endogenously expressed FUS, TDP-43, or hnRNPA1 in HEK293T cells.** HEK293T NTC and DNAJB6 KO cells were transfected with an EYFP expression vector and endogenous levels of TDP-43, FUS, and hnRNPA1 were assessed.

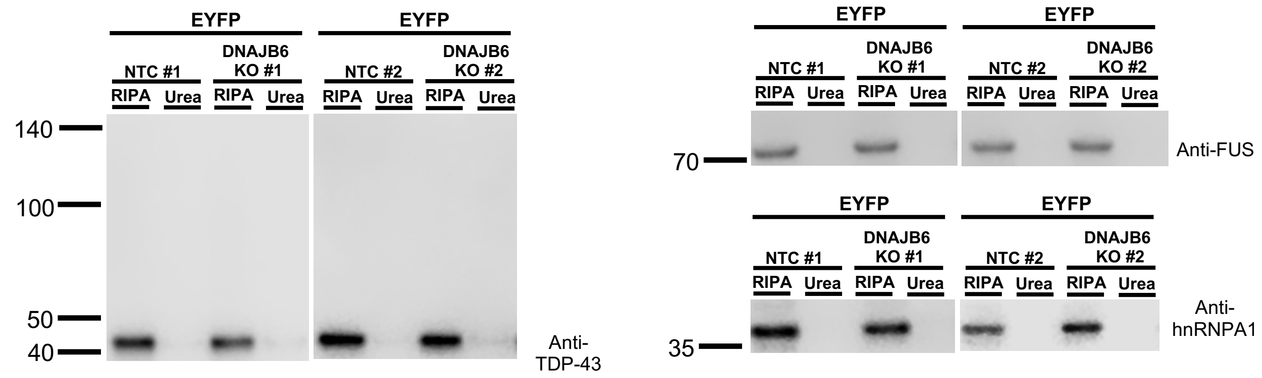

**Supplementary Figure 15. Biophysical characterization of DNAJB6 with clients**. **a**. DNAJB6 at 3 µM concentration undergoes LLPS at physiological salt concentrations, sample imaged 30 minutes after dilution. **b.** Ability of AF555 labeled DNAJB6 at 0.25 µM to LLPS in 500 mM NaCl and “physiologic” 50 mM NaCl conditions. Samples were imaged at 30 minutes. Scale bar represents 5 microns. **c.** FUS-mEmerald and AF555 labeled DNAJB6 co-mingle when mixed at physiological salt concentrations and at an endogenous (6:1) ratio, 1.5 µM and 0.25 µM, respectively. Scale bar represents 10 microns. Samples were imaged 20 minutes after mixing. **d.** FUS-mEmerald (1.5 µM) alone and FUS-mEmerald + DNAJB6 (1.5 µM + 0.25 µM) condensates were subjected to FRAP 30 minutes after condensate formation.

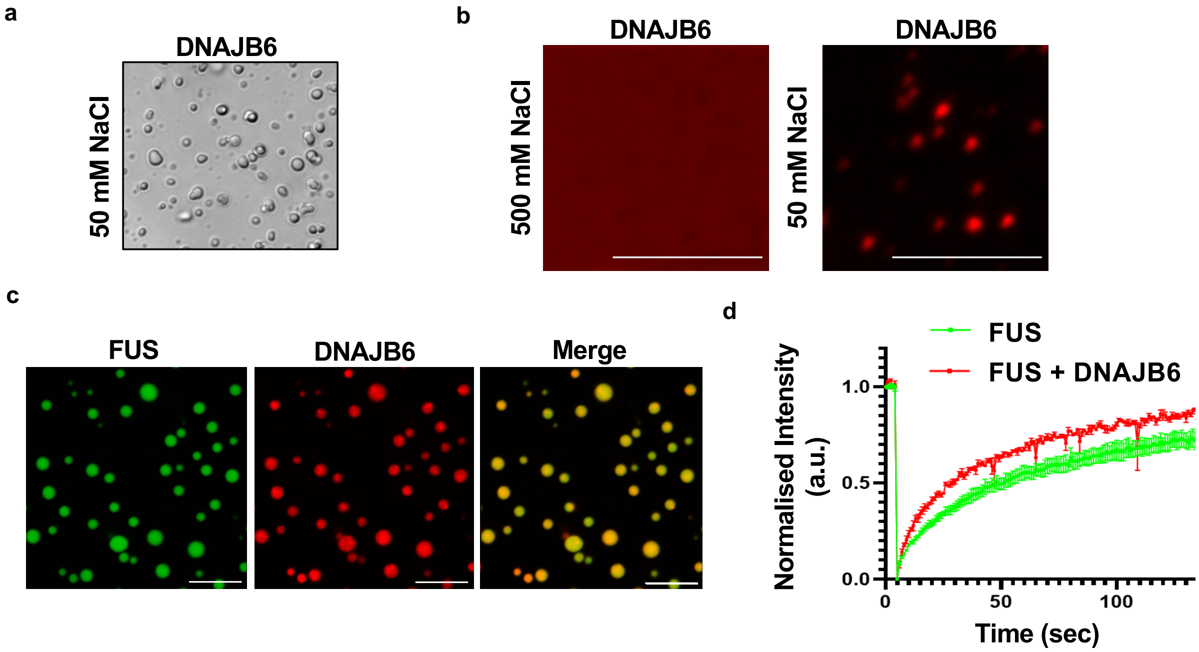

**Supplementary Figure 16. Example of AFM-IR nano-chemical analysis on single FUS+DNAJB6 condensates.** Maps of **a.** 3-D morphology, **b.** IR absorption in the Amide I (1655cm-1), **c.** Contact resonance by phase locked loop (PLL). **d.** IR spectra from 10 independent locations (each location 5 co-averaged spectra) on the condensate, **e.** their average + SE and **f.** second derivative of the amide I band to deconvolve protein secondary structure contributions. **g.** 3-D morphology maps of 3 independent FUS and FUS + DNAJB6 condensates

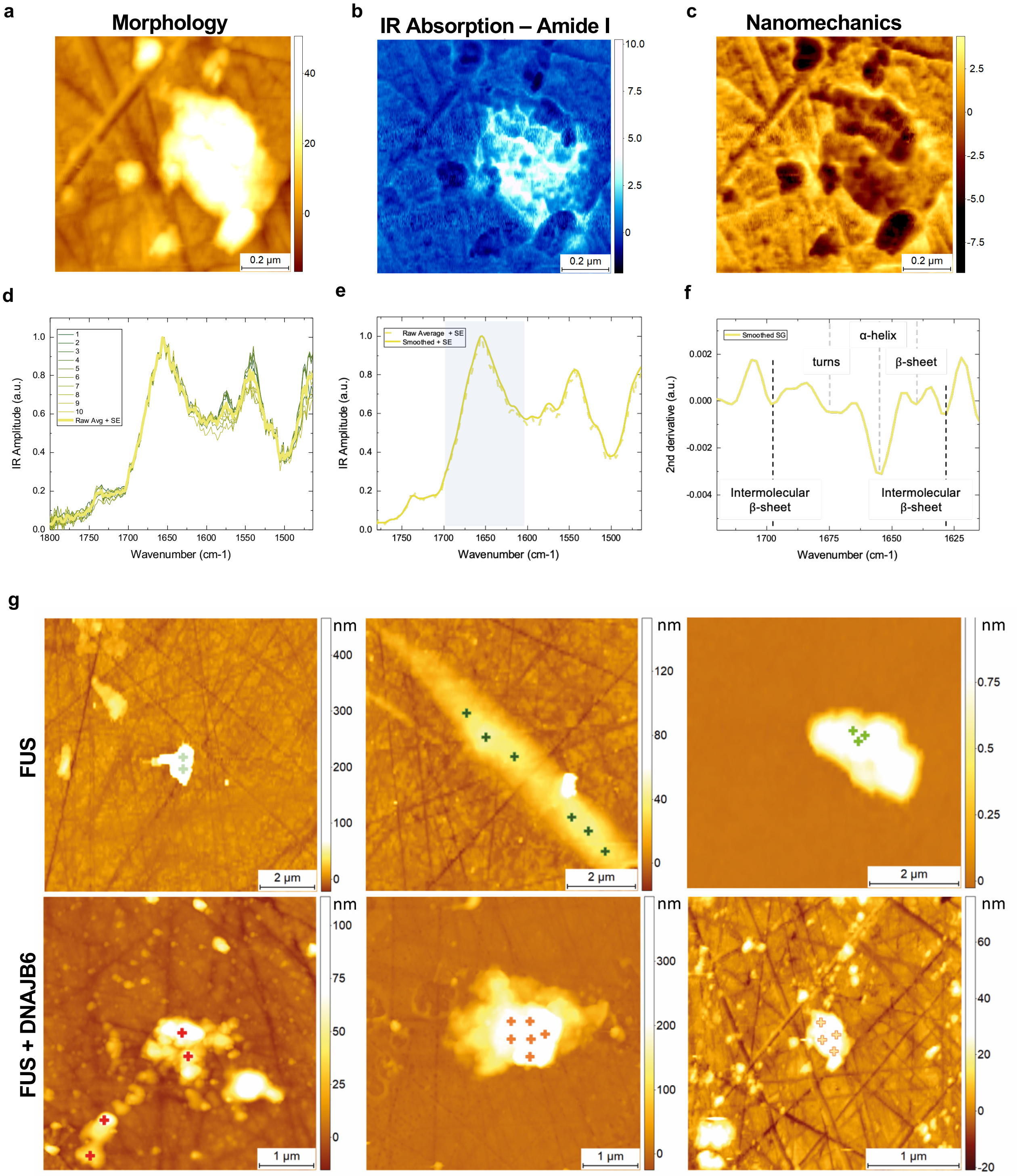

**Supplementary Figure 17. Identification of DNAJB6 domains important for activity in yeast. a.** Testing domain deletions and J domain H31Q loss of function point mutant for their ability to rescue the FUS expressing yeast model. ΔJ, ΔG/F, ΔS represent deletion of the J-domain, glycine-phenylalanine rich, or serine rich region of DNAJB6, respectively. **b.** Expression confirmation of DNAJB6 and mutant variants. Comparisons were conducted with ordinary one-way ANOVA; ns = not significant, ***P≤0.001, ****P≤0.0001.

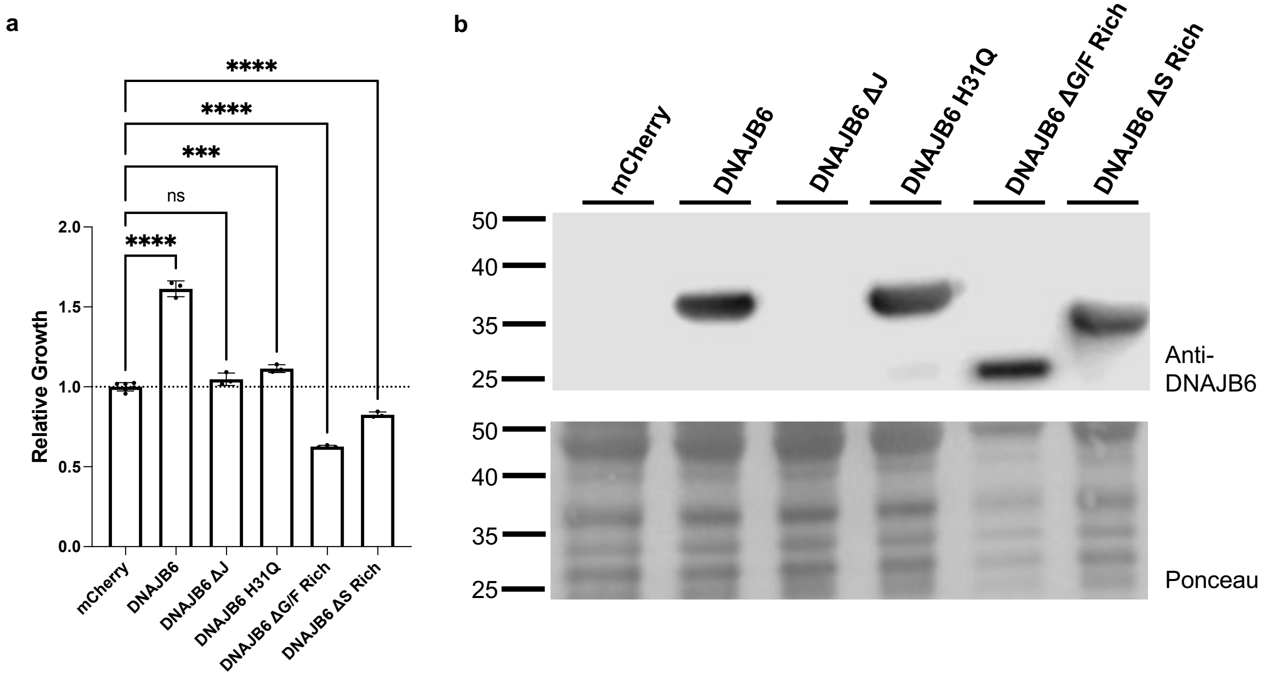

**Supplementary Figure 18. Correlation statistics between biological replicates of deep mutational scan of DNAJB6. a.** Correlation between log2 fold changes for all amino acid changes at all positions tested in the deep mutational scanning approach. **b.** Correlation between log2 fold changes for amino acids in each set of 14 amino acids analyzed together in one sequencing batch.

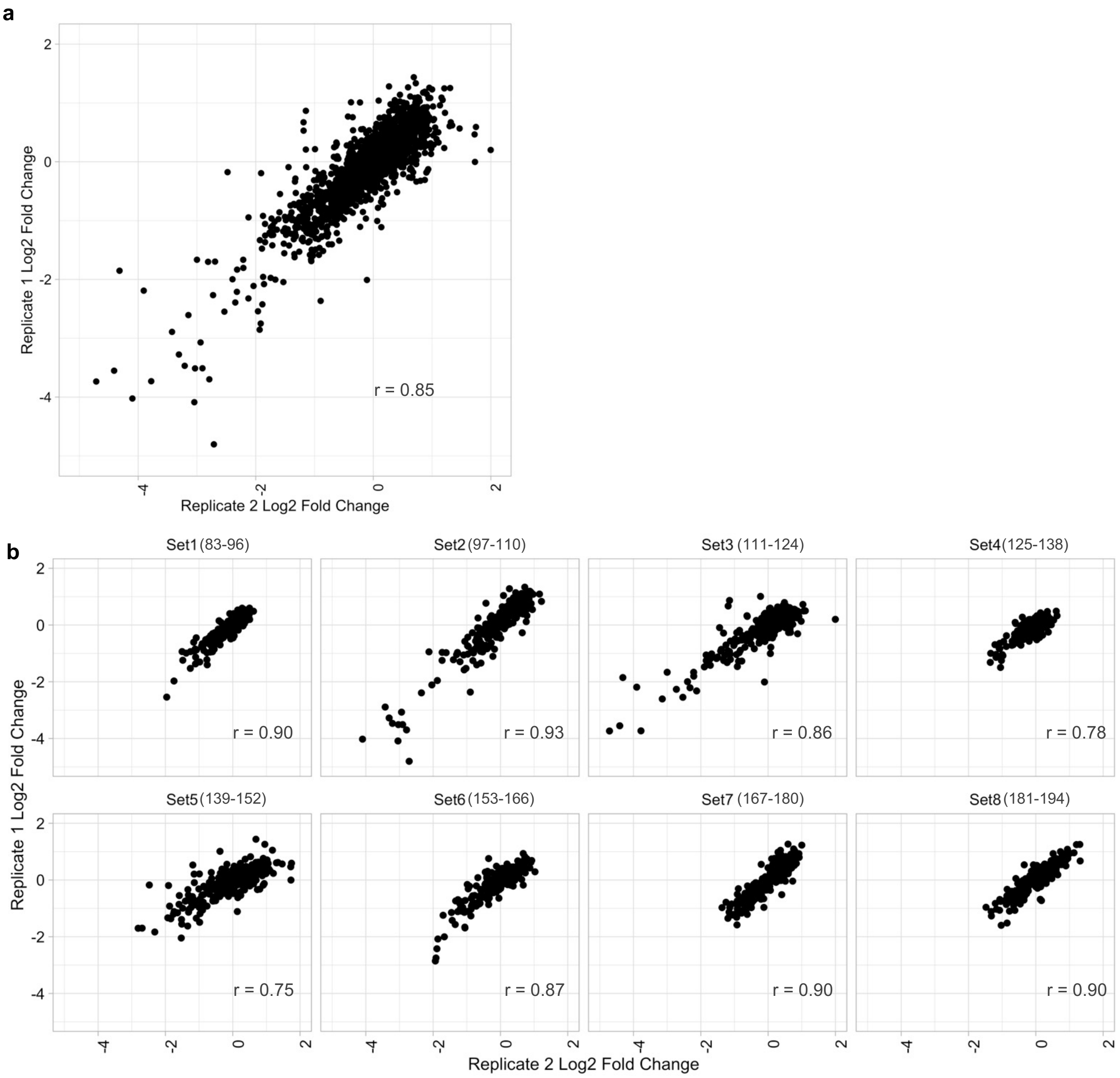

**Supplementary Figure 19. Increasing amount of FUS plasmid transfected increases the amount of SDS-insoluble, Urea-soluble species.** Different doses of FUS expression plasmid were transfected and protein was harvested 48 h after transfection to assess the solubility of FUS.

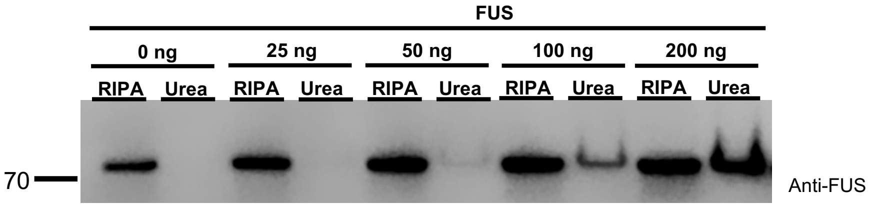

### Supplementary Tables

**Supplementary Table 1. Models included in pool.** Names, sequences, and 20 bp barcodes of models in the pool. Growth and passaging conditions for each model's secondary validation are also included.

**Supplementary Table 2. Results of chaperone screen.** Primary data from analysis pipeline for each interaction in the chaperone screen data set. Log2FC = Log2 fold change, FDR = False Discovery Rate.

**Supplementary Table 3. Results of secondary validation of called hits and suspected interactions from the chaperone screen.** Three replicates for each condition were tested.

**Supplementary Table 4. Results of ORFeome screen.** Primary data from analysis pipeline for each interaction in the chaperone screen data set. Log2FC = Log2 fold change, FDR = False Discovery Rate.

**Supplementary Table 5. Results of secondary validation of called hits and suspected interactions from the orfeome screen.** Three replicates for each condition were tested.

**Supplementary Note 1**

To begin to establish the feasibility of multiplex high-throughput screening, we first needed to determine the reproducibility of all the required steps within the pipeline. Towards this goal, we assembled a pilot pool composed of 117 DNA-barcoded yeast strains. Among the barcoded strains in the pool were several proteotoxic models with known genetic rescuers such as yeast prions RNQ1 and SUP35, and NDD models such as FUS, TDP-43, and alpha-synuclein. Also included in this pilot pool were other proteotoxic models selected to represent a range of different strengths of toxicity to assess how variation in the amount of growth arrest caused by a model (i.e. mild, moderate, and strong), affects the reproducibility of our system. Taking advantage of the scalability of the DNA-barcoding and to help control for variation at the biological and technical levels, each model was transformed into 3 unique isogenic DNA-barcoded strains (i.e. redundantly barcoded). This allows each barcoded variant of the same model to serve as an "internal biological replicate", and for the collective behavior of all barcodes associated with the same model to be used to determine the effects of each tested genetic modifier.

The first set of experiments that were performed was testing whether *en masse* mating and selection of the pilot DNA-barcoded pool was consistent when performed across multiple wells each mated to the same control rescuer strain. We observed strong correlation between separately mated pools, suggesting relative barcode abundance is preserved through mating and selection (Sup. Fig. 2a-b). We next determined whether individually mated and selected diploid pools resulted in a reproducible behavior for all members of the library when inoculated into inducing media and allowed to grow back to saturation. We observed strong correlation between separately mated, selected, and outgrown pools. These data suggest that the pool shows a consistent behavior across replicate experiments and that endpoint measurements of barcode abundance can be used to make comparisons between control rescuer and active rescuer wells (Sup. Fig. 2c-d).

**Supplementary Note 2**

Using the pilot pool, we tested whether altering the relative abundance of particular strains in the pool might improve our ability to detect known, literature-reported interactions between molecular chaperones and the library of proteotoxic models [27,78,79]. The "All" pooling strategy evenly mixed all 117 strains. The "Low" pooling strategy evenly mixed all strains but excluded a number of control yeast strains expressing proteins that lack toxicity (e.g. enhanced yellow fluorescent protein) to enable more division opportunities before the pool reached growth saturation. The "Skew" pooling strategy mixed all 117 strains but seeded strong and moderately toxic models at a higher initial abundance compared to the mild and non-toxic models. We observed comparable performance between the Low and All pools, with 16/17 and 14/17 literature-reported positive controls demonstrating positive log2 fold change when the behavior of all the DNA-barcodes associated with the same model were averaged (Sup. Fig. 3a). In sharp contrast, the Skew pool showed the worse performance detecting only 12/17 positive control interactions, along with showing overall lower log2 fold changes as compared to the Low and All pools.

Upon further examination of the resulting data, a stronger correlation between biological replicates using the Low pooling strategy mated to the same benign rescuer over other strategies was observed (Sup. Fig. 3b). Additional analysis was performed in which the relationship between the coefficient of variation (CV) of a barcode and its mean relative abundance in the pool was examined. As previously shown in both RNA-sequencing and microbiome sequencing datasets, low abundance members in a mixed pool tend to show higher variance in their abundance values, which we hypothesize may render more toxic models within the pool (which are rapidly depleted during outgrowth) more variable [80,81]. The ability of a pooling strategy to reduce variability at all sampling levels, in particular those with lower abundance, suggests it should have improved performance and increased sensitivity to detect real interactions. The Low pooling strategy was generally associated with lower variability for barcodes at all relative abundances. The Skew strategy did reduce the variability of lowly abundant barcodes primarily associated with highly toxic models compared to the All pooling strategy, but was also associated with higher variability for less toxic, generally more abundant models possibly as a result of their lower initial seeding (Sup. Fig. 3c).

Taking the Low pooling strategy forward, we assessed sources of biological and technical noise. For this study, we considered a biological replicate to require a separate mating, selection, outgrowth, DNA harvest, and PCR amplification for sequencing. We considered technical replicates to be separate PCR amplification reactions performed on the same harvested DNA for sequencing. We observed relatively minor sources of both biological and technical variation (Sup. Fig. 4a). We also assessed whether averaging between multiple biological or technical replicates improved the reproducibility of the screen by reducing the CV ~ relative abundance relationship of barcoded strains. Averaging relative abundances of barcode strains between multiple replicates reduced the variability of barcoded strains, with averaging between 2 replicates conferring a similar advantage to averaging between 3 replicates (Sup. Fig. 4b-c). This suggested that a screening paradigm that adopts the Low pooling strategy with two biological replicates for each well and two technical replicates for each biological replicate is optimized for sensitive detection of genetic modifiers of proteotoxicity. We tested whether the Low pooling strategy along with two biological or technical replicates improved performance over the initial pilot experiment and observed that log2 fold changes were stronger and captured all known interactions (Sup. Fig. 5a). We validated, via spot assay, all potential interactions within this pilot interaction space and observed strong concordance with screen data (Sup. Fig. 5b-g)

**Supplementary Note 3**

The initial variance modeling data suggested that lower abundance members of the pool are highly variable and would restrict the assay to detecting only strong interactions for these models. We hypothesized that merging information between isogenic "redundantly barcoded" strains would help improve the detection of mild and moderate interactions for lower abundance pool members. Using the prior association of CV ~ relative abundance, we modeled the necessary fold change in order to detect statistically significant enrichment of models while also accounting for the large degree of multiple hypothesis testing when implementing the approach. To model the required fold change to detect significant interactions at an $\alpha = 0.05$ with a pool of 50 models, we determined the multiple hypothesis corrected Z-score necessary to reach significance with a Bonferroni correction. From this Z-score, we derived the necessary fold change required to reach significance from the CV at each relative abundance. To model sharing information between barcodes, we used Stouffer's Z-score method to simulate the required individual Z-scores necessary for significance when these Z-scores are combined. Without information sharing between isogenic redundantly barcoded strains containing the same model, greater than 2-fold change in abundance was necessary for significance for lowly abundant barcodes. Our modeling suggested that information sharing between isogenic strains representing the same model would enable more sensitive detection of weaker interactions, similar to how information is shared between multiple gRNAs in CRISPR screens to identify essential genes [76]. By pooling information between 5 or more isogenic strains, we determined that rescuers that increased the abundance of a model within the mixed pool by 1.5 fold could be detected with statistical significance (Sup. Fig. 6). With this approach, we observed that the benefits of redundant barcoding scaled faster than the penalties of multiple testing, suggesting that additional redundant barcoding is favorable for sensitive detection of interactions.

For each model, we assembled 5-7 individual barcoded strains and validated equal growth between isogenic strains containing the same model (Sup. Table 1). We pooled a total of 302 barcoded strains and assessed the CV ~ relative abundance relationship with this new pool to determine the proper read depth. We observed a similar CV ~ relative abundance relationship between this larger pool and the pilot pool used to optimize our approach. We hypothesized that increasing the read depth of each well may also reduce the CV of lowly sampled barcodes. However, we observed similar CV ~ relative abundance profiles, with 128,000 or greater reads per well demonstrating the minimum read depth required to gain most of the benefits of increased read depth in terms of CV ~ relative abundance and number of lowly sampled barcodes that are captured (Sup. Fig. 6a-b). This suggests that lowly abundant barcoded strains may retain inherent variance as a result of the degree of proteotoxicity and growth suppression they experience. At this level of sequencing depth, 24 96-well plates can be sequenced on a single Illumina NextSeq 75bp High Output run, with an approximate cost of $0.70 per screened well.
